# Topography of corticopontine projections is controlled by postmitotic expression of the area-mapping gene Nr2f1

**DOI:** 10.1101/2021.05.10.443413

**Authors:** Chiara Tocco, Martin Øvsthus, Jan G. Bjaalie, Trygve B. Leergaard, Michèle Studer

## Abstract

Axonal projections from layer V neurons of distinct neocortical areas are topographically organized into discrete clusters within the pontine nuclei during the establishment of voluntary movements. However, the molecular determinants controlling corticopontine connectivity are insufficiently understood. Here, we show that an intrinsic cortical genetic program driven by Nr2f1 graded expression is directly implicated in the organization of corticopontine topographic mapping. Transgenic mice lacking cortical expression of *Nr2f1* and exhibiting areal organization defects were used as model systems to investigate the arrangement of corticopontine projections. Combining three-dimensional digital brain atlas tools, *Cre*-dependent mouse lines, and axonal tracing, we show that *Nr2f1* expression in postmitotic neurons spatially and temporally controls somatosensory topographic projections, whereas expression in progenitor cells influences the ratio between corticopontine and corticospinal fibres passing the pontine nuclei. We conclude that cortical gradients of area patterning genes are directly implicated in the establishment of a topographic somatotopic mapping from the cortex onto pontine nuclei.

**Summary statement:** Cortical gradient expression of the area patterning gene *Nr2f1* spatially and temporally controls corticopontine topographic connectivity in layer V projection neurons.

## Introduction

Neuronal populations responsible for fine motor coordination are arranged in topographically organized maps in the neocortex and cerebellum, with different body parts being represented in largely continuous maps in the somatosensory cortex (Chapin and Lin, 1984; Fabri and Burton, 1991; Welker, 1971; Woolsey and Van der Loos, 1970), and discontinuous, fractured maps in the cerebellum (Bower, 2011; Bower et al., 1981; Bower and Kassel, 1990; Leergaard et al., 2006; Nitschke et al., 1996; Shambes et al., 1978). The intercalated regions of this network, the pontine nuclei, deep cerebellar nuclei, and thalamus, receive and integrate signals ultimately resulting in coordinated and seamlessly executed behaviours (Buckner, 2013; Peterburs and Desmond, 2016; Stoodley and Schmahmann, 2010), including fine voluntary movements (Badura et al., 2013; Mottolese et al., 2013).

The pontine nuclei constitute the major synaptic relay for cerebro-cerebellar signals (Brodal and Bjaalie, 1992; Lemon, 2008; Mihailoff et al., 1985). Axonal projections originating from layer V pyramidal neurons across the neocortex are distributed in topographically organized clusters within the pontine nuclei, as shown in monkey (Brodal, 1978; Schmahmann and Pandya, 1997), cat (Bjaalie and Brodal, 1997), rat (Leergaard et al., 2000a; Leergaard et al., 2000b), and to some extent also in mice (Henschke and Pakan, 2020; Inoue et al., 1991; Proville et al., 2014). Within the pontine nuclei, the three-dimensional (3D) arrangement of clustered terminal fields, well described in rats, both preserves the overall topographical relationships of the cortical maps, but also partially overlap and introduce new spatial proximities among projections from different cortical areas (Bjaalie and Brodal, 1989; Leergaard, 2003; Leergaard and Bjaalie, 2007).

To date, the mechanisms responsible for establishing the topographic map between the neocortex and pontine nuclei are poorly understood. The leading proposition, referred to as chrono-architectonic hypothesis, postulates that the complex 3D topography is a product of straightforward spatial-temporal gradients, possibly combined with non-specific chemo-attractive mechanisms (Leergaard, 2003; Leergaard and Bjaalie, 2007; Leergaard et al., 1995). Recent new discoveries open the possibility that other mechanisms are also in action during the establishment of the corticopontine maps. Several lines of evidence point to a functional role of gradients in gene expression during topography of sensory maps in several systems (D’Elia and Dasen, 2018; Erzurumlu et al., 2010; Fritzsch et al., 2019; McLaughlin and O’Leary, 2005), but whether this process is also operative during establishment of corticopontine topography is not understood. A recent study has shown that postmitotic graded expression of the HOX gene *Hoxa5* is directly involved in imparting an anterior to posterior identity to pontine neurons and attract corticopontine axons (Maheshwari et al., 2020). Whether gradient expression of molecular factors along the antero-posterior or medio-lateral axes of the cerebral cortex also intrinsically determine the topography of corticopontine projections is still not known. Layer V neurons from the anterolateral cerebral cortex project to the central regions of the pontine nuclei, while more medially located cortical regions project to more external parts; motor area projections are distributed more medially and rostrally, while somatosensory ones reach the middle and caudal parts of the pontine nuclei. Finally, auditory and visual cortical projections innervate the dorsolateral regions of the pontine nuclei (Leergaard et al., 2004; Leergaard and Bjaalie, 2007). The fine-tuned and precise corticopontine topography leaves open the possibility for cortical neurons being intrinsically programmed to target specific groups of pontine neurons, possibly coupling both intrinsic and extrinsic mechanisms in directing proper topographical innervation to the pontine nuclei.

A new theme of cortical patterning has emerged, in which genetic factors direct the spatial and temporal establishment of topographically organized axonal connections between the cortex and subcortical brain regions (Cadwell et al., 2019). Area mapping genes are expressed in gradients along the different axes of the cortical primordium, are known to modulate the size and position of future cortical areas (Alfano and Studer, 2012; Cadwell et al., 2019; O’Leary and Sahara, 2008), and to determine areal fate and regulate expression of downstream molecules that in turn control the topographic organization of synaptic inputs and outputs of related structures (Assimacopoulos et al., 2012; Greig et al., 2013). These genes represent good candidates for modulating topographic mapping. In mice, the *Nr2f1* gradient expression appears to be a particularly strong candidate for having a formative role during the establishment of topographic maps (Armentano et al., 2007; Liu et al., 2000; Zhou et al., 2001). For instance, *Nr2f1* is expressed in cortical progenitor cells from embryonic day E9.0 in a high caudo-lateral (future sensory) to low rostro-medial (future motor) gradient fashion, which is maintained in postmitotic descendants and postnatally when the cortical area map is completed (Bertacchi et al., 2019; Flore et al., 2017; Tomassy et al., 2010). Previous studies show that Nr2f1 promotes somatosensory (S1) by repressing motor identity in postmitotic neurons and in its absence area size and thalamocortical topography are both affected (Alfano et al., 2014; Armentano et al., 2007; Bertacchi et al., 2019). We thus hypothesized that Nr2f1 could represent one of these factors able to control topographic corticopontine mapping during corticogenesis.

To this purpose, we made use of cortico-specific *Nr2f1* conditional knockout mice as an in *vivo* model system and a paradigm to investigate the contribution of cortical genetic programs in the establishment of topographic corticopontine projections. Two distinct conditional mouse lines were crossed to the *Thy1-eYFP-H* reporter line (Feng et al., 2000), in which YFP is highly expressed in cortical layer V pyramidal neurons and their axonal projections (Porrero et al., 2010). The distribution of fluorescent YFP signals as well as anterogradely labelled corticopontine projections were evaluated by comparison of spatially corresponding microscopic images and 3D visualization of extracted point-coordinate data representing labelling. Our results indicate that cortical Nr2f1 expression plays a dual role in controlling the spatial-temporal development of corticopontine projections. While early expression in progenitor cells influences the ratio between corticofugal fibres passing the pontine nuclei, loss of postmitotic late expression specifically affects topographic pontine mapping. Overall, our data demonstrate that intrinsic genetic programs and postmitotic graded expression of cortical area mapping genes are implicated in the establishment of area-specific targeting of corticopontine neurons.

## Results

### Benchmark 3D topographic organization of corticopontine projections in wild-type mice

To first establish a 3D reference of the topographical organization of corticopontine projections in wild-type adult mice, we used tract tracing data from the *Allen Institute Mouse Brain Connectivity Atlas* (Wang et al., 2020), see flowchart in **Figure 1A**. These data allowed us to visualize the spatial distribution of the pontine projections of motor and somatosensory neocortical areas. Corticopontine projections digitized from sagittally-oriented microscopic images (matching the orientation used for our experimental data, see below) were co-visualized as 3D data points in a 3D viewer tool (see e.g. **Figure 2A-D**). Experimental data were selected and coloured according to tracer injection localizations in the cerebral cortex, to first visualize and compare the pontine distribution of corticopontine projections from motor and somatosensory areas, and secondly to determine the organization of projections arising from progressively more medial and caudal locations in the cerebral cortex, following the cortical neurogenetic gradient that ripples out from the anterolateral cortex (Leergaard and Bjaalie, 2007; Smart, 1984). Our findings confirm that the somatosensory and motor neurons of the mouse cortex project to largely separate parts of the pontine nuclei (Henschke and Pakan, 2020; Inoue et al., 1991; Proville et al., 2014), with clustered terminal fields topographically distributed in the same concentric fashion (**Figure 2L-P**) as previously reported in rats (Leergaard et al., 2000a; Leergaard and Bjaalie, 2007). These 3D point data were also used below as supplementary control data, and as benchmarks for interpreting YFP expression and tract-tracing results in *Nr2f1* mutant mice.

**Figure 1.**
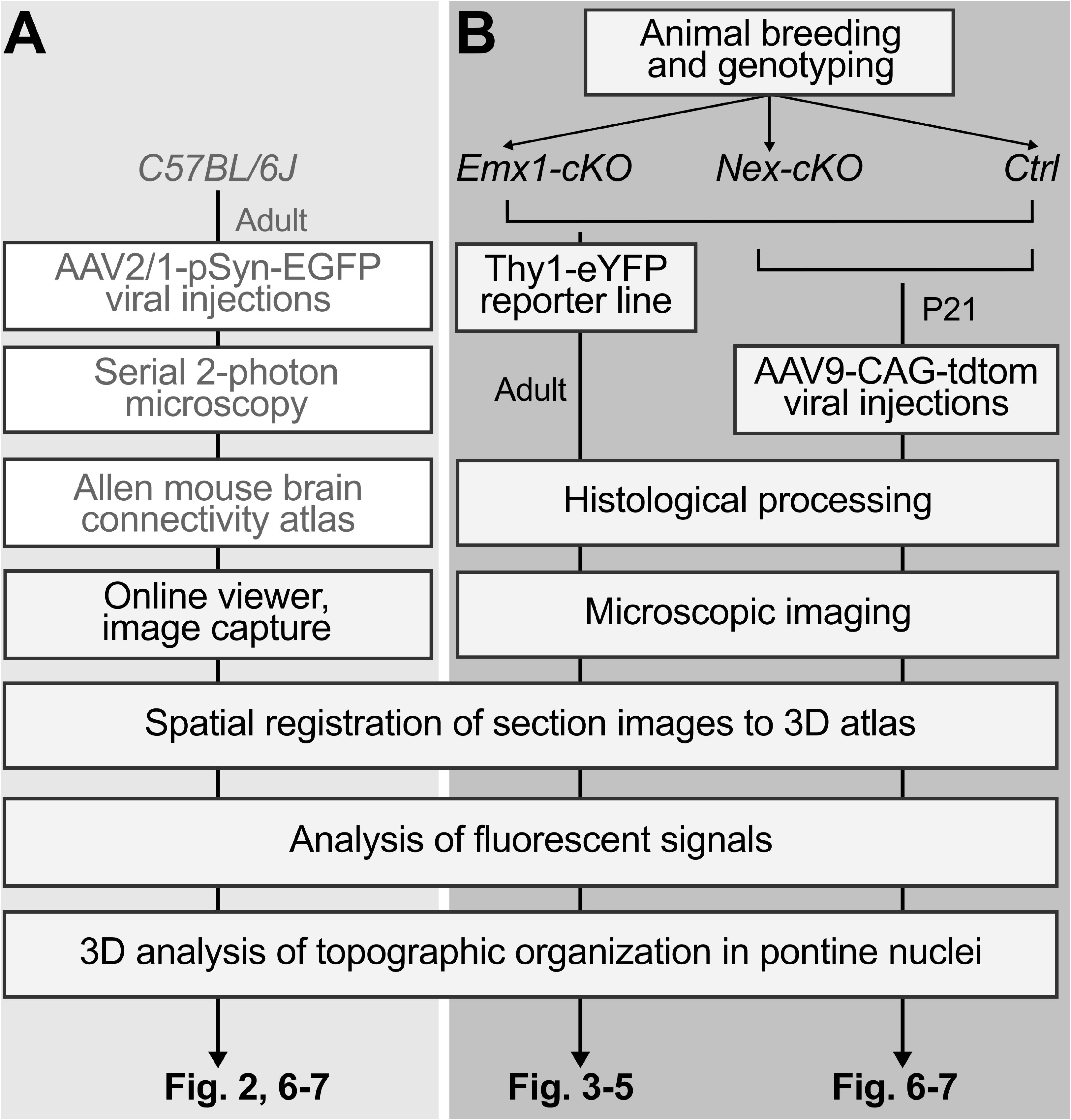
Experimental and analytic workflow. (**A**) Workflow for generating a 3D topographic map of corticopontine projections in wild-type mice using tract tracing data from the Allen Mouse Brain Connectivity Atlas, mapped and compared in a 3D reference atlas space. Steps performed by the Allen institute are indicated with light grey text in white boxes. (**B**) Middle and right columns represent the two paradigms investigated in cKO models, with the analytic steps performed in adult control, Emx1-cKO and Nex-cKO mutant animals (middle column), and the tract tracing of the 3D topography of motor and somatosensory corticopontine projections in young control and Nex-cKO mutants (right column). All images were spatially registered to the Allen mouse brain atlas (CCFv3; Wang et al., 2020) prior to analyses, to facilitate comparison of images and spatial distribution patterns. Results are shown in Figures 2-7.

**Figure 2.**
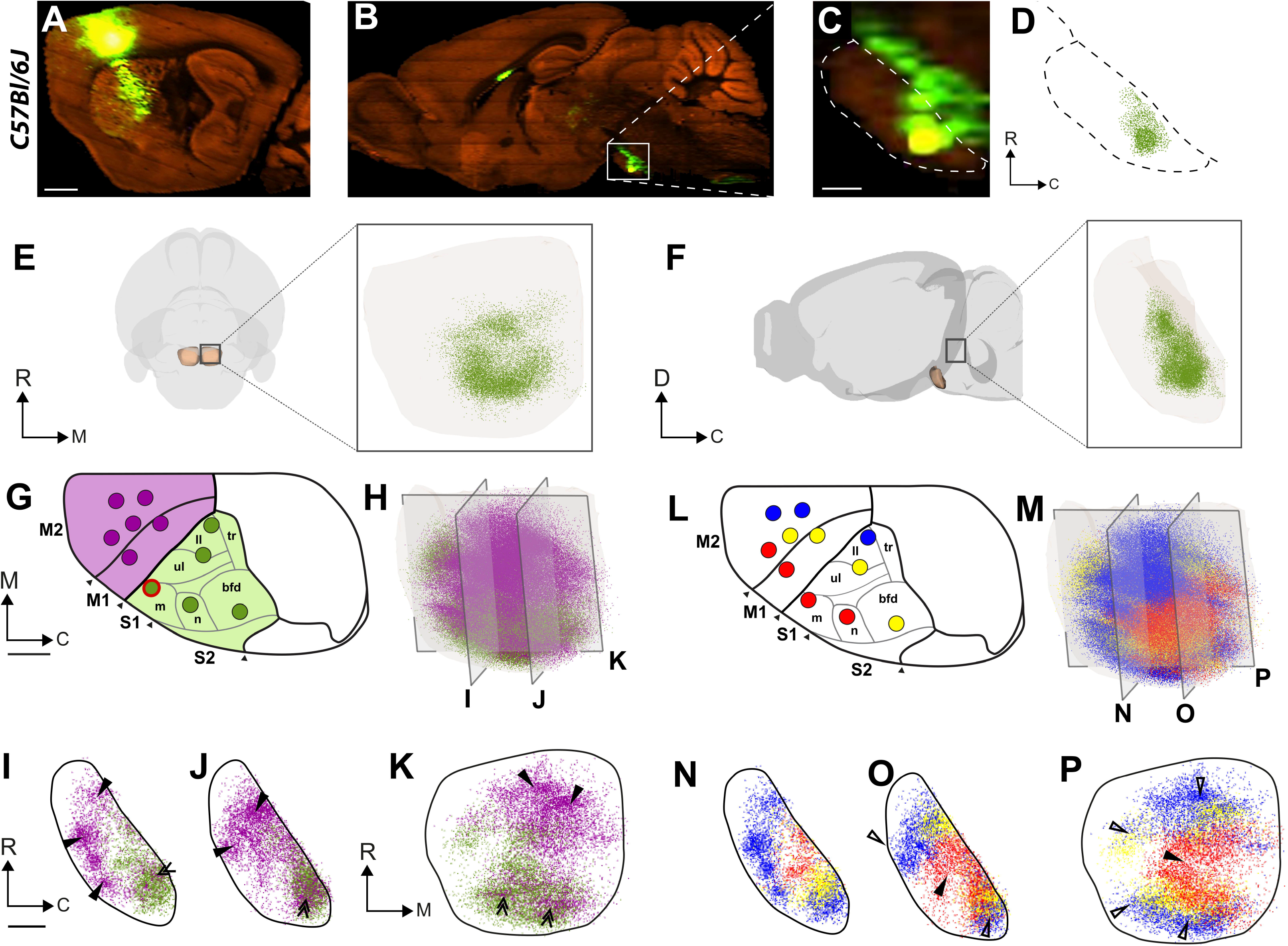
Topographical organization of corticopontine projections in wild-type mice. (**A-D**) Representative example illustrating the data acquisition of corticopontine projections labelled by viral tracer injection in the S1 face representation of a wild-type mouse from the Allen Mouse Brain Connectivity Atlas. The tracer injection site centre (**A**) and anterogradely labelled axons in the pontine nuclei (**B, C**) are shown in sagittal microscopic images. (**D**) Dot map representation showing semi-quantitatively recorded points corresponding to the density of labelling observed in the section shown in (**C**). Panels **E,F** show the 3D point populations recorded from the example case, together with a transparent surface rendering of the right pontine nuclei, seen in view from ventral (**E**) and medial (**F**). The S1 corticopontine projections are distributed in dense clusters located centrally in the pontine nuclei. (**G-K** and **L-P**) Differently coloured 3D visualizations of point clouds representing spatial distribution of anterogradely labelled corticopontine axons derived from 11 experiments available in the Allen Mouse Brain Connectivity Atlas, injected with the anterograde tracer EGFP in the primary (M1)/secondary (M2) motor cortex (purple dots) or primary somatosensory (S1) cortex (green dots), at locations indicated with colour coded circles in **G** and **L**. (**H, M**) 3D visualizations of all points in the right pontine nuclei, colour coded as indicated in **G** and **L**. A grid of transparent grey planes indicate the position and orientation of ~100 μm thick digital slices cut with sagittal and frontal orientations through the point clouds, shown in **I-K**, and **N-P**. The red circle in G indicates the tracer injection site shown in **A**. (**G-K**) Point clouds showing that motor and somatosensory areas largely target different parts of the pontine nuclei, with projections from M1 and M2 (purple dots) located more peripherally towards rostral, ventral, and medial than projections from S1 (green dots) as indicated by arrowheads in I-K, but also that motor and sensory projections overlap caudally in the pontine nuclei (double arrowheads in **I-K**). (**L-P**) 3D co-visualization of all data points colour-coded in red, yellow or blue according to the location of the cortical injection sites from anterolateral (red) progressively towards medial or posterior (yellow, blue). The slices through the point clouds reveal a concentric arrangement in the pontine nuclei, with projections from the anterolateral parts of the M1/M2 and S1 located centrally and medially (arrowheads in **O-P**), and projections from more medial and posterior cortical locations progressively shifted towards rostral, caudal, and lateral (unfilled arrowheads in (**O-P**). Abbreviations: bfd, barrel field; C, caudal; D, dorsal; ll, lower limb; m, mouth; M1, primary motor cortex; M2, secondary motor cortex; n, nose; R, rostral; S1, primary somatosensory cortex; S2, secondary somatosensory cortex, tr, trunk; ul, upper limb; V, ventral. Scale bars, 1 mm (**A, B**), 200 μm (**C,I**).

### Area-specific layer V neuron distribution in cortices lacking Nr2f1

To assess the influence of cortical area mapping on the establishment of topographical organization in mouse corticopontine projections, we used *Nr2f1 conditional knock-out* mice as an experimental model system and their littermates as controls (Alfano et al., 2014; Armentano et al., 2007). Two well-established conditional *Nr2f1* mouse mutants were used: the *Nr2f1^fl/fl^::Emx1-Cre* mouse, in which *Nr2f1* expression is abolished from early cortical progenitor cells at mouse embryonic (E) age 9.5 (Armentano et al., 2007), and the *Nr2f1^fl/fl^::Nex-Cre* mouse in which *Nr2f1* expression is inactivated at later stages (E11.5-E12), solely in cortical postmitotic neurons (Alfano et al., 2014; Goebbels et al., 2006). Both mouse lines were crossed to the *Thy1-eYFP-H* reporter line to specifically restrict signal expression to layer V pyramidal neurons (Harb et al., 2016; Porrero et al., 2010). In these mice, YFP expression follows the physiological distribution of subcortical layer V projections, including corticospinal and corticopontine fibres, with a high density in motor cortex and a gradually decreased distribution in S1 and more caudal areas. For simplicity, the lines are below named *Emx1-cKO* and *Nex-cKO*, respectively.

In agreement with a previous study (Porrero et al., 2010), we observed substantial YFP signal expression in the hippocampus, tectum, and pontine nuclei, as well as in the globus pallidus, claustrum, endopiriform nucleus, nucleus of the lateral olfactory tract, mammillary nuclei, piriform area, and the substantia innominata in adult mice (**Figure 3A**). Signal expression was also seen in the vestibular nuclei, deep cerebellar nuclei, and cerebellum. Although YFP expression was present in almost the same cortical regions in 2-months-old mutant mice as in controls, detailed analysis of layer V expression revealed some distinct differences in the spatial distribution of *Emx1-cKO* and *Nex-cKO* brains relative to their respective controls, and between the two conditional lines. We used an ImageJ macro to automatically count YFP+ nuclei by area of interest in the brain using a threshold based on intensity and shape of the elements. This allowed to estimate the number of YFP+ neurons in seven cortical areas (corresponding to prefrontal, motor, somatosensory, auditory, visual and retrosplenial cortices in control mice) defined by delineations from spatially-registered overlay images from the *Allen Mouse Brain Atlas* (Wang et al., 2020); **Figure 1; Figure 3B-D’**). In control animals, the number of YFP+ neurons follows the physiological distribution of subcortical layer V projections (**Figure 3B, B’**), as previously reported (Polleux et al., 1997; Porrero et al., 2010; Shepherd, 2009). Strong staining in motor and somatosensory cortices resulted in bright signal expression in the cerebral peduncle (CP) and CST (**Figure 3A**).

**Figure 3 -.**
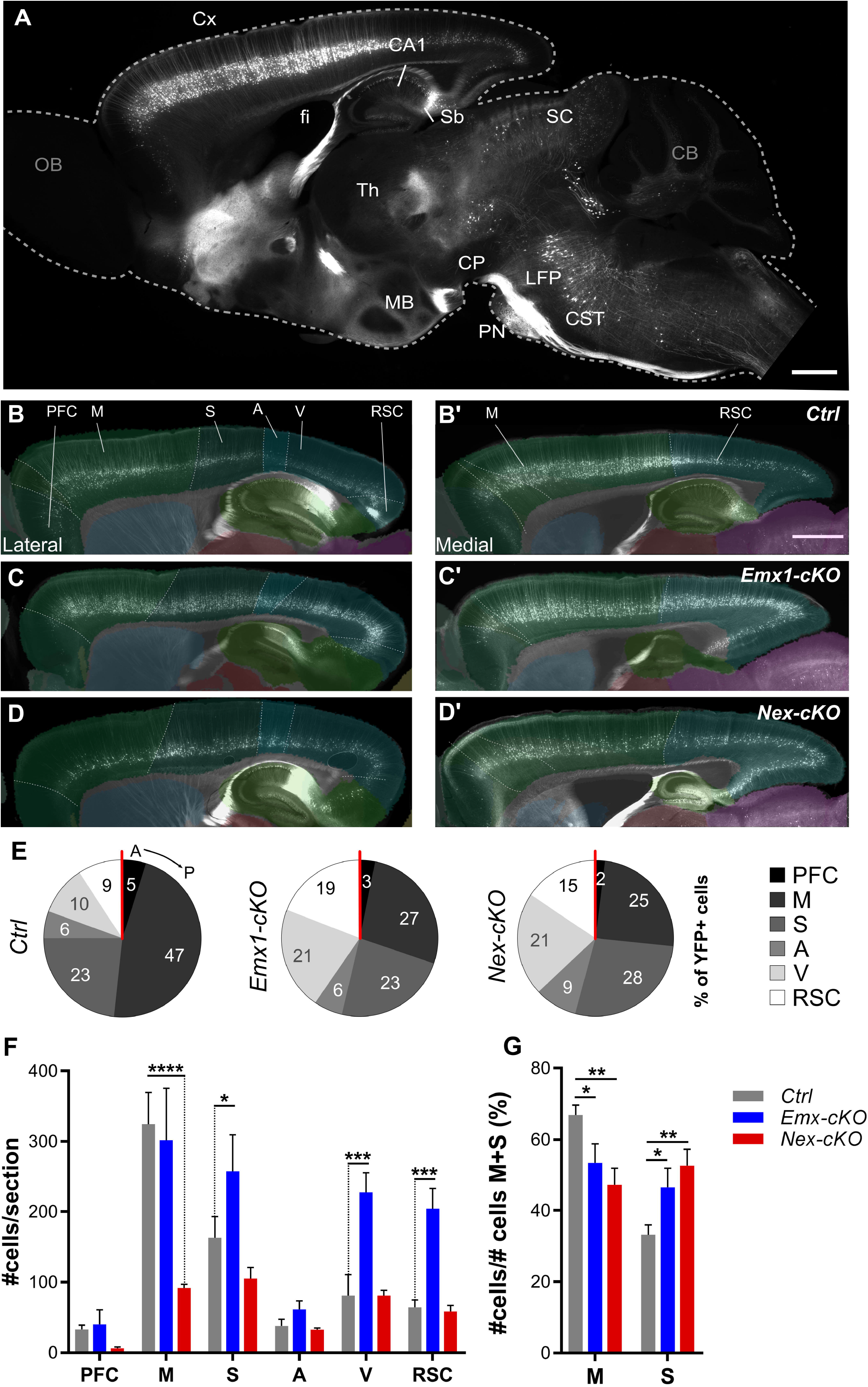
Distribution of YFP-positive layer V pyramidal neurons in cerebral cortex of control, Emx1-cKO and Nex-cKO adult brains. (**A**) Fluorescence microscopy image of a representative sagittal section from control brain, showing widespread YFP expression. (**B-D’**) Fluorescence microscopy images of laterally (**B-D**) and medially (**B’-D’**) sagittal sections in Ctrl (**B,B’**), Emx1-cKO (**C,C’**), and Nex-cKO (**D,D’**) brains, with spatially corresponding CCFv3 atlas diagrams superimposed to indicate the location of cortical areas. (**E**) Pie charts illustrating the distribution of YFP+ layer V neurons in different cortical areas along the antero-posterior axis of controls and mutant brains. (**F**) Column graph showing the number of YFP+ neurons across cortical areas in adult Ctrl (grey), Emx1-cKO (blue) and Nex-cKO (red) mice normalized for the total sections analyzed. (**G**) Column graph showing a subsampling of the analysis in **E**. Here, the number of YFP+ layer V neurons in frontal (motor; M) and parietal (somatosensory; S) areas are normalized for the sum of YFP+ cells of M and S regions only, and the values are presented as percentages. Data were analyzed with 2way-ANOVA test (see also **Supplementary Table 4**), and are illustrated as mean ± SEM (Ctrl, n=6; Emx1-cKO, n=4, Nex-cKO, n=4). Abbreviations: A, auditory cortex; CA1, cornu ammonis area 1; CB, cerebellum; CP, cerebral peduncle; CST, corticospinal tract; Cx, cortex; fi, fimbria; GC, gustatory cortex; LFP, longitudinal fascicle of the pons; M, motor areas (includes primary and secondary motor cortices); MB, mammillary body; OB, olfactory bulb; PFC, prefrontal cortex; PN, pontine nuclei; RSC, retrosplenial cortex; S somatosensory areas; SC, superior colliculus; Sb, subiculum; Th, thalamus; V, visual cortex. Scale bars, 1mm (**A**); 500μm (**B-D; B’-D’**).

To understand whether the spatial organization of YFP+ layer V cortical neurons was affected upon loss of cortical *Nr2f1* gradient expression, we quantified and compared the number of YFP+ cells across mutant and control brains (**Figure 3E, F**). Since previous reports showed that the distribution of layer V neurons changes in the absence of Nr2f1 (Alfano et al., 2014c; Armentano et al., 2007a; Tomassy et al., 2010), we expected that the Thy1-YFP signal would also be altered, as a read-out of layer V changes. Indeed, the layer V gradient was disrupted in both mutant strains, and YFP+ cells were more homogenously distributed along the anteroposterior cortical axis (**Figures 3B-F**). While *Emx1-cKO* brains showed a significant increase of YFP+ cells in parietal and occipital regions, compared to controls, this was less the case for *Nex-cKO* brains, which instead revealed a decrease only in frontal areas (**Figure 3F**). To better quantify the differences between corticopontine projections from frontal and parietal areas, where normally the motor and somatosensory areas develop (Leergaard et al., 2000a; Leergaard et al., 2000b), we compared the ratio of YFP+ cells over the total cell populations counted in the frontal (motor) and parietal (somatosensory) areas (**Figure 3G**). We found that the percentage of YFP+ cells was significantly decreased in frontal (motor) and concomitantly increased in parietal (somatosensory) in both *Emx1-cKO* and *Nex-cKO* cortices compared to controls, but no differences were found between the two lines (Figure 3E, G). This lack of difference in the distribution of YFP+ layer V neurons between cortical regions normally containing the motor and somatosensory areas, is in line with the acquisition of a motor-like identity in somatosensory cortex of Nr2f1-deficient brains, as previously reported (Alfano et al., 2014). We next asked whether and how alterations in layer V organization observed in the two mutant lines would be translated into layer V corticospinal projections and/or corticopontine topographic mapping.

### Abnormal corticospinal projections and fasciculation in *Nr2f1* mutant brains

We hypothesized that the disordered cortical distribution of YFP-expressing layer V neurons in mutant mice might influence the integrity of subcortical axonal projections. In all cases, strong YFP signal expression was seen bilaterally in the main corticofugal pathways (**Figure 3A**), visible as longitudinally oriented fibre bundles coursing towards the CP (**Figure 4A-D**), passing dorsal to the pontine nuclei as the longitudinal fasciculus (**Figure 4B’-D’**), and continuing through the brain stem towards the spinal cord as the CST. Since a large fraction of the corticobulbar fibres terminate in the pontine nuclei (Tomasch, 1968; Tomasch, 1969), we reasoned that abnormal distribution of YFP+ layer V neurons observed in mutant mice (**Figure 3**) might affect corticopontine innervation, and could be reflected in an abnormal size of the pontine longitudinal fascicle as it enters the CP and exits the pons in rostral and caudal positions to the pontine nuclei, respectively. To evaluate this, we measured the dorsoventral width of the longitudinal fascicle of the pons in sequential sections along the medio-lateral axis. The measurements were taken at rostral and caudal levels to the pontine nuclei (**Figure 4A’**). Surprisingly, we found the lateral part of the fascicle to be wider at both rostral and caudal levels in *Nex-cKO* mice compared to *Emx1-cKO* and controls, while being narrower medially (see red area chart in **Figure 4E, F**), suggesting that the longitudinal fascicle of the pons is flattened and expanded laterally upon Nr2f1 inactivation in postmitotic neurons. Only minor differences were observed in the *Emx1-cKO* fascicles (blue area chart in **Figure 4E, F**). This is also supported by quantification of the total surface of the longitudinal fascicle of the pons at rostral and caudal levels, which shows a significant surface reduction at caudal, but only a tendency at rostral levels in *Emx1-cKO* mice (**Figure 4G, H**). These data indicate that loss of Nr2f1 in cortical progenitors, but not in neurons, results in fewer YFP+ fibres passing the pontine nuclei towards the brain stem to form the CST.

**Figure 4 -.**
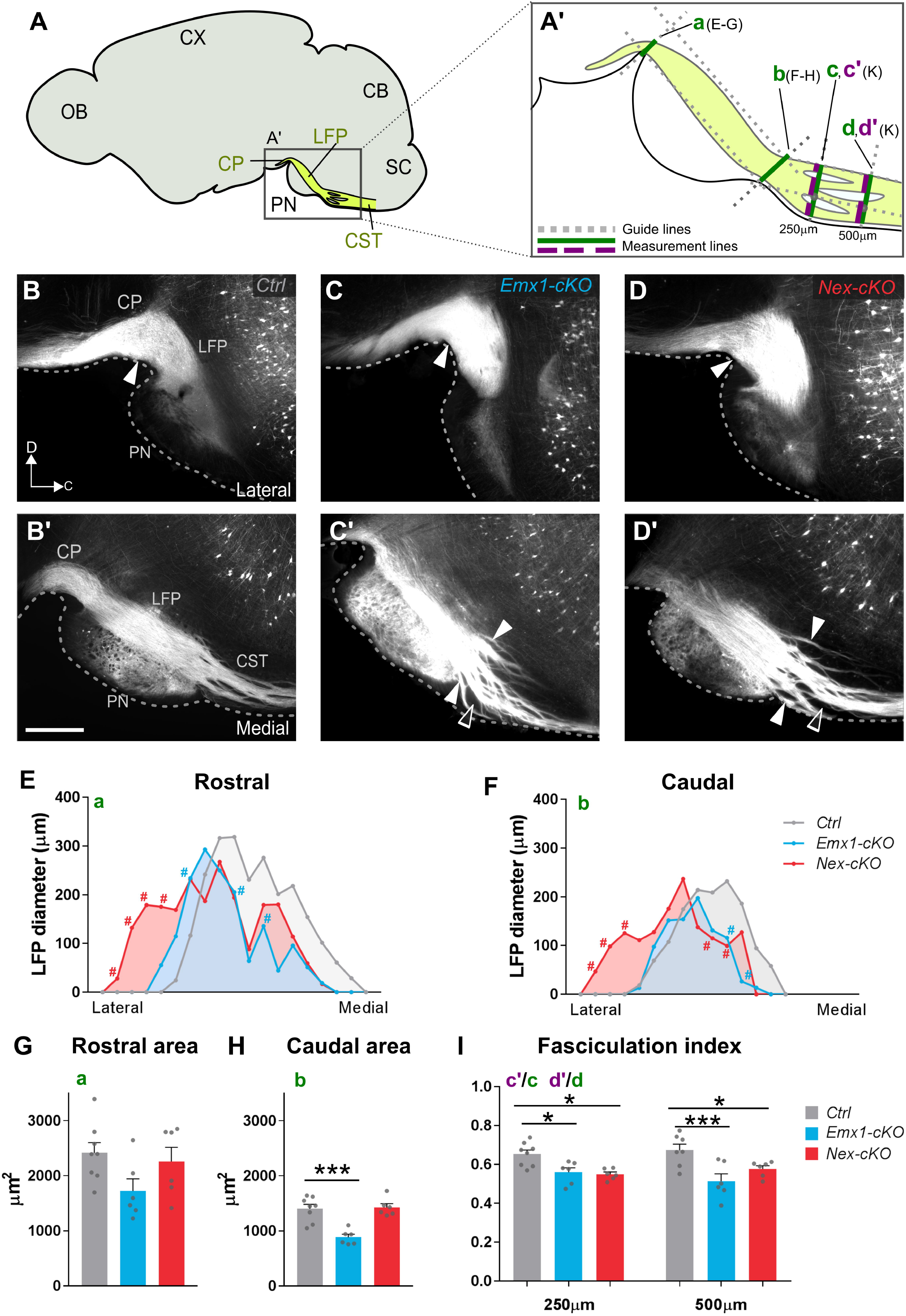
Loss of Nr2f1 function leads to abnormal corticospinal projections and fasciculation. (**A**) Schematic diagram of a sagittal mouse brain section showing the location of the pontine nuclei (PN) and descending fibre tracts (yellow) in the cerebral peduncle (CP), longitudinal fascicle of the pons (LFP) and corticospinal tract (CST). (A’) Diagram illustrating the different measurements shown in E-I. (**B-D** and **B’-D’**) Fluorescence microscopy images of sagittal sections showing the corticospinal tract entering the LFP (**B-D**) and continuing caudal to the pons as the CST. White arrowheads point to the sites of measurement plotted in **E** and **F**, respectively. Rostrally, the labelled LFP has similar thickness in the three genotypes. Caudally, the CST features defasciculation of fibre bundles (full arrowheads in **C’,D’** point to thinner and more dispersed bundles in Emx1-cKO and Nex-cKO mutants), empty arrowheads point to empty spaces between bundles. (**E, F**) Plots showing LFP diameter measurements obtained from lateral to medial before and after innervating the PN (rostral and caudal respectively). Each measurement represents the average value of corresponding sections among distinct animals and each position on the x-axis represents a specific section of the series. (**G-H**) Column graphs showing average values of the area under the curves in **E-F**. A comparable number of fibres reach the CP in the three genotypes (**G**). In Emx1-cKO brains fewer fibres are seen to exit the level of the pons compared to control- and Nex-cKO brains (**H**). (**I**) Column graph showing CST fasciculation index, based on measurements of total thickness and fibre thickness (green and purple line respectively in **A’**) performed at 250 and 500 μm from the terminal edge of the PN. A ratio between the two measurements was calculated for each position. Data are represented as mean ± SEM. Data were analysed with 2way-ANOVA test (**E-F**) or ordinary one-way ANOVA test (**G-I**) and corrected for multiple comparison with the Bonferroni test (see also Supplementary Table 4). Ctrl, n=8; Emx1-cKO, n=6; Nex-cKO, n=6. #<0.05 (**E-F**) * < 0.05, **< 0.01, ***<0.005 (**G-I**). Abbreviations: CB, cerebellum; CX, cortex; OB, olfactory bulb; SC, spinal cord. Scale bar, 500μm (**B’**).

Moreover, we observed, caudal to the pontine nuclei, abnormally widespread fibre fascicles in the CST of mutant animals (arrowheads in **Figure 4C’, D’**). To determine whether this was a significant difference between animal groups, we estimated the degree of fibre bundle fasciculation in the CST of *Emx1-cKO* and *Nex-cKO* mice. At locations of 250μm and 500μm caudal to the pontine nuclei (**Figure 4A’**), we measured the total dorsoventral width of the CST at several medio-lateral levels and subtracted the gaps between the YFP-expressing fibre bundles at the same levels. The ratio of the total width of the CST and fibres was used as a measure of the fasciculation index (**Figure 4I**). Notably, in both groups of mutant mice, we found a lower degree of fasciculation in the CST which was more pronounced at the most caudal level (**Figure 4I**). Together, these data show that loss of *Nr2f1* expression affects the diameter, shape, and degree of fasciculation of the CST originating from layer V neurons.

### Dual role of Nr2f1 in targeting corticopontine projections

Next, we evaluated whether topographical organization of corticopontine projections dependents on proper cortical area mapping and layer V distribution (**Figure 5**). To this purpose, we assessed the spatial distribution of YFP signal expression within the pontine nuclei by comparing intensity-normalized microscopic images of spatially corresponding sagittal sections from the brains of *Emx1-cKO, Nex-cKO* and control littermate animals (**Figure 1B; Figure 5A**). A complete documentation of spatially comparable and reproducible images is provided in **Supplementary Figures 1 and 2**. In control brains, we observed a strong YFP signal in central parts of the pontine nuclei, with the densest expression tending to surround a centrally located zone exhibiting less dense signals (**Figure 5B-E**). This region of the pontine nuclei typically receives strong projections from somatosensory areas (green dots in **Figure 2G-K**). Some signal expression was also visible in medial parts of the pontine nuclei (**Figure 5D, E**), which is known to primarily receive projections from the cortical motor areas (purple dots in **Figure 2G-K**). By contrast, signal expression was lower in rostral and lateral parts of the pontine nuclei (**Figure 5B, C, E**), which are known to receive projections from visual and auditory areas of the cerebral cortex (Inoue et al., 1991; Leergaard and Bjaalie, 2007).

**Figure 5 -.**
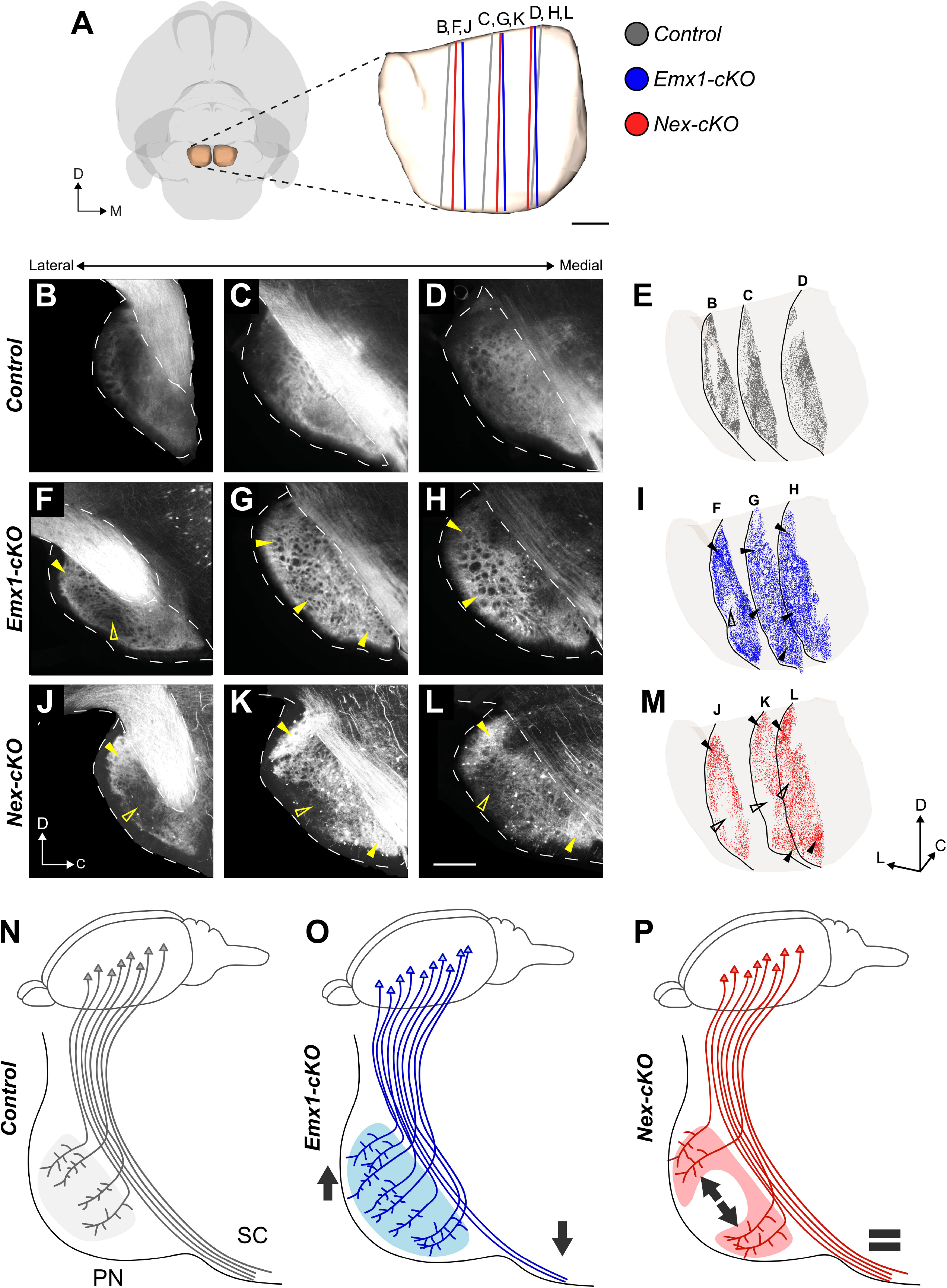
Distribution of YFP signal expression in the pontine nuclei of knock-out and control mice. (**A**) 3D representation of the outer surface of the brain (transparent grey) and pontine nuclei (PN) (transparent brown) in view from ventral. Coloured lines indicate the location and orientation of the sagittal sections shown in **B-M**. (**B-D, F-H, J-L**) Fluorescence microscopy images of sagittal sections from corresponding medio-lateral levels of the PN, showing the spatial distribution of YFP signal expression in control, Emx1-cKO, and Nex-cKO mice, respectively. (**E, I, M**) 3D visualization of the PN in an oblique view from ventro-medial, with point coded representations of signal expression from each of the three sagittal sections shown in **B-D, F-H**, and **J-L**, respectively. Filled yellow or black arrowheads point to regions with increased signal expression in mutant mice, while non-filled arrowheads indicate regions with decreased signal expression. In control mice (**B-E**), signal expression is primarily seen in central and caudal parts of the pontine nuclei, while in Emx1-cKO mice (**F-I**) signal expression is more widespread and diffuse throughout the entire PN, including more peripheral parts towards rostral, ventral and caudal positions (filled arrowheads in **G-I**). In Nex-cKO mice (**J-M**), signal expression is reduced in the central core region of the PN (non-filled arrowheads in **K-M**), while being increased in peripheral (rostral and caudal) regions. (**N-P**) Diagrams summarizing observed changes in corticopontine connectivity upon Nr2f1 inactivation. In control mice (**N**), YFP^+^ corticopontine projections primarily target central part of the PN and a substantial number of fibres continue towards the spinal cord. In Emx1-cKO mutants (**O**), fewer fibres reach the SC and more projections target the PN (down- and up-pointing grey arrows, respectively) and show a pattern of innervation more diffused and widespread that in controls. In Nex-cKO animals (**P**), no difference between corticospinal and corticopontine projections are detected compared to controls (grey equal sign), but corticopontine topography appears to be affected, whereby fibres reach more lateral, motor-receiving PN regions instead of targeting the S1-receiving core (illustrated by grey divergent arrows). Abbreviations: C, caudal; D, dorsal; L, lateral; M, medial, PN, R, rostral; SC, spinal cord. Scale bar, 200 μm.

Interestingly, *Emx1-cKO* mice showed a relatively homogeneous signal distribution across all parts of the pontine nuclei, and displayed more expression in the dorsolateral regions (**Figure 5F-I**). Signal expression was also present in the medial part of the nuclei (**Figure 5G-I**). This observation fits well with the finding of a higher number of YFP+ neurons in the occipital cortex (**Figure 3C, C’, F**), which projects to the dorsolateral pontine nuclei. By comparison, *Nex-cKO* animals showed more constrained signal expression and predominated in rostrally and caudally located clusters extending from the cerebral peduncle towards the ventral surface of the pons, and medially surrounding a central core with low expression (**Figure 5J-M**). These clusters were more peripherally located than the ones observed in control animals (**Figure 5J-M**). Notably, little signal expression was seen in the central region of the PN of all *Nex-cKO* cases (unfilled arrowheads in **Figure 5J-L**), despite the presence of YFP+ layer V neurons in somatosensory cortex (**Figure 3D, D’, F, G**). This central region is normally innervated by projections from the face representations located in **S1** (**Figure 2A-F**).

Taken together, these findings show that corticopontine projections are abnormally distributed in *Nr2f1* cortical deficient mice, with more homogenously (non-specific) distributed expression in *Emx1-cKO* mice (schematically summarized in **Figure 5N,0**), and more peripherally distributed signal expression in *Nex-cKO* mice, that display reduced expression in the central region of the pontine nuclei normally receiving somatosensory projections (schematically summarized in **Figure 5N,P; Supplementary Figures 1 and 2**). In both mutant groups, the signal expression was expanded to dorsolateral regions of the pontine nuclei that normally are innervated by projections from occipital cortical areas. This suggests that cortical Nr2f1 graded expression in postmitotic neurons might be directly involved in the establishment of topographically organized corticopontine projections.

### Altered somatosensory topographic projections in *Nex-cKO* adult mutant mice

To further support that *Nr2f1* gradient cortical expression might be directly involved in topographical pontine mapping, we decided to focus on the *Nex-cKO* genetic model and unveil the cortical origin of the innervation defect. Compared to the *Emx1-cKO* mice, in which projections appear more disorganized and abundant, we considered the *Nex-cKO* mouse model, exhibiting externally shifted projections, more suitable for investigating the changes in topographical mapping using experimental tract tracing techniques. We injected the AAV9-CAGtdTomato anterograde viral tracer (Pourchet et al., 2021) in the cortex of 5 days-old (P5) *Nex-cKO* mice and littermate controls, in frontal and parietal locations corresponding to the motor or S1 areas in control mice, respectively (**Figures 6, 7**). The mice were sacrificed at P21 and brain sections analysed microscopically (**Figure 1B**). All histological sections were spatially registered to the *Allen Mouse Brain Atlas* (common coordinate framework, CCF3; (Wang et al., 2020)), and the location of tracer injections sites were mapped in the same atlas space (**Figures 6A, 7A**). For each injection site location in a *Nex-cKO* brain, we selected the most corresponding control experiment or wild-type tract-tracing data from the *Allen Mouse Brain Connectivity Atlas* as additional controls. Representative examples of injections and comparisons between controls and littermate *Nex-cKO* brains are shown in **Figures 6B-C’ and 7B-C’**. Complete documentation of all tracing experiments cases is provided in ***Supplementary Figures 3-5***.

**Figure 6.**
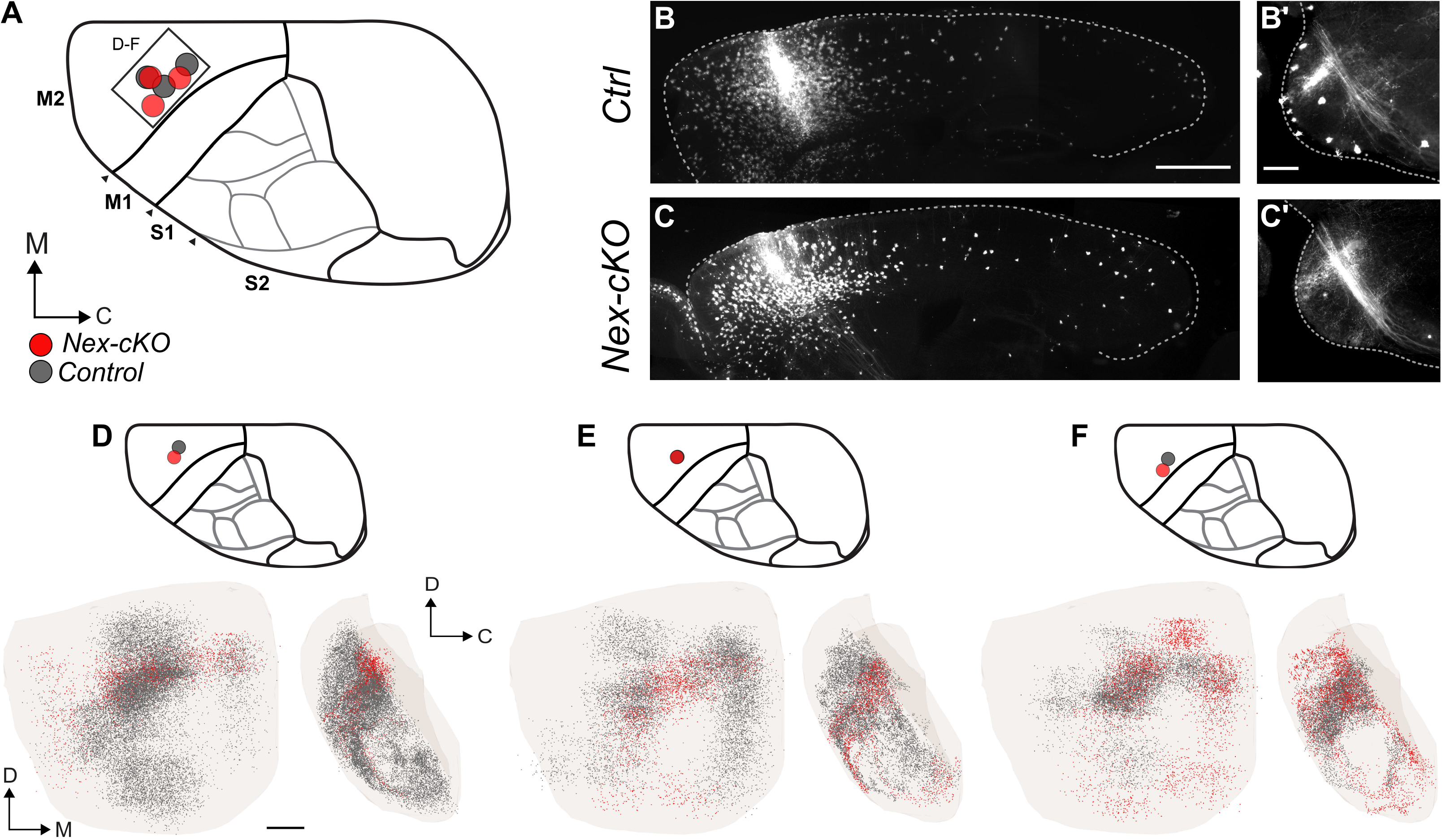
Anterograde tracing of corticopontine projections from frontal (motor) areas in Nex-cKO mice. (**A**) Overview of injection sites in corresponding locations in the right frontal cortex in Nex-cKO (red) and control (grey) brains. The atlas diagrams shown in **A, D, E**, and **F** indicate the control cortical area identities. (**B, B’** and **C, C’**) Representative microscopic images of the injection site localization (**B-C**) and pontine nuclei innervation (**B’, C’**) in the control and Nex-cKO cases reported in panel **F**. (**D-F**) 3D coloured point clouds representing axonal labelling in corresponding pairs of Nex-cKO (red) or control/wild-type (dark grey) mice, shown within a transparent surface representation of the right pontine nuclei in ventral and medial views. Inset drawings of the brains seen from dorsal show the location of tracer injection sites for each combination of point clouds. Tracer injections in corresponding locations in frontal cortex of both Nex-cKO and control/wild-type mice give rise to similar corticopontine labelling in rostrally located clusters, curving towards ventral and caudal along the surface of the pontine nuclei. Abbreviations: C, caudal; D, dorsal; M, medial; M1, primary motor cortex; M2, secondary motor cortex; S1, primary somatosensory cortex; S2, secondary somatosensory cortex. Scale bars, 1mm (**B**) 200 μm (**B’,D**).

**Figure 7.**
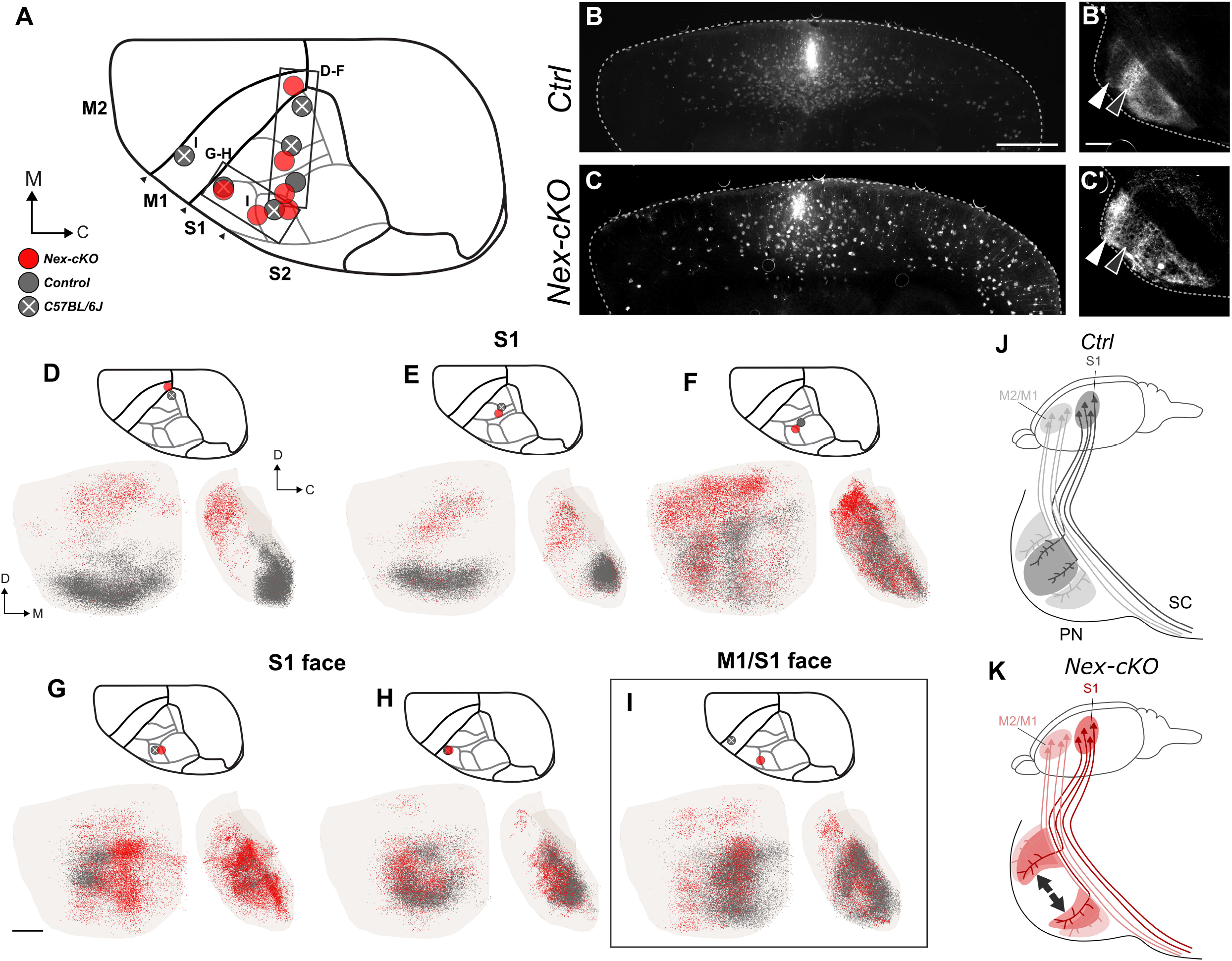
Anterograde tracing of corticopontine projections from somatosensory areas in Nex-cKO mice. (**A**) Overview of injection sites in corresponding locations in the right parietal (primary somatosensory; S1) cortex in Nex-cKO (red), control (grey), and wild-type (grey with white cross) brains. (**B, B’** and **C, C’**) Representative microscopic images of the injection site localization (**B-C**) and pontine nuclei innervation (**B’, C’**) in the control and Nex-cKO cases reported in panel **F**. (**D-I**) 3D coloured point clouds representing axonal labelling in corresponding pairs of Nex-cKO (red) or control/wild-type (grey) mice, shown within a transparent surface representation of the right pontine nuclei in ventral and medial views. Inset drawings of the brains seen from dorsal show the location of tracer injection sites for each combination of point clouds. Corresponding tracer injections in corresponding locations in the medial parietal cortex of Nex-cKO and control/wild-type mice give rise to labelling in different parts of the pontine nuclei, with corticopontine projections in control mice distributed in elongated curved clusters located caudally (grey points in **D,E**) or laterally in the pontine nuclei (grey points in **F**), while projections from the same locations in Nex-cKO mice are shifted to more peripheral rostral and lateral parts of the pontine nuclei (red points in **D-F**). All tracer injections in anterolateral part of the parietal cortex of Nex-cKO and control mice, in regions corresponding to the S1 face representation in wild-type mice, gave rise to labelling in the central region of the pontine nuclei, however with a subtle medial shift of projections in Nex-cKO brains (**G**, see also **I**). Corresponding tracer injections in the most anterolateral part of S1 in a Nex-cKO and control gave rise to highly similar labelling, centrally in the pontine nuclei. (**I**) Tracer injections in widely separated locations in S1 (Nex-cKO) and M1 (wild-type control) gave rise to largely corresponding labelling in the medial part of the central core region of the pontine nuclei, albeit with additional rostral and medial labelling in the Nex-cKO experiment. (**J-K**) Diagrams summarizing observed changes in corticopontine connectivity upon Nr2f1 postmitotic inactivation. In control mice (**J**), projections from motor areas (M2/M1, pink) and S1 (purple) target largely segregated parts of the pontine nuclei, with somatosensory projections targeting the central core region, while motor projections target more peripheral rostral, caudal, and medial parts of the pontine nuclei. In Nex-cKO animals, corticopontine topography of S1 is affected, whereby fibres reach lateral, motor-receiving PN regions, instead of targeting the core (illustrated by grey divergent arrows). Abbreviations: C, caudal; D, dorsal; M, medial; M1, primary motor cortex; M2, secondary motor cortex; PN, pontine nuclei; S1, primary somatosensory cortex; S2, secondary somatosensory cortex, SC, spinal cord. Scale bars, 1mm (**B**), 200 μm (**G**).

In controls, the spatial distributions of corticopontine projections (dark grey point clouds in **Figures 6 and 7**) were highly comparable with the labelling patterns seen in corresponding wild-type tracing data from the *Allen Mouse Brain Connectivity Atlas*. As expected, tracer injections into areas corresponding to the motor cortex in control mice gave rise to labelled axonal clusters located rostrally, caudally, and medially in the pontine nuclei (purple dots in **Figure 2G-K, Figure 6D-F; Supplementary Figures 3 and 4**). In all *Nex-cKO* mice receiving tracer injections into areas normally corresponding to the motor cortex, the overall distribution of corticopontine labelling was found to be essentially similar to that observed in control cases (compare grey point with red point clouds in **Figure 6D-F; Supplementary Figures 3-5**).

Tracer injections into S1 areas of control animals gave rise to labelled axonal clusters located centrally or caudally in the pontine nuclei (grey point clouds in **Figure 7D-I**). The spatial distribution of labelling varied systematically as a function of the location of the cortical injection sites, such that the 1) face representation in the lateral part of S1 projects centrally in the pontine nuclei (**Figure 7G,H; Supplementary Figure 3I,J**), 2) the whisker representations lateral and more posterior in S1 project to regions surrounding the central core (**Figure 7F; Supplementary Figure 3K**), while 3) the more medially located forelimb and hindlimb representations of S1 projected to medio-laterally oriented regions in the caudal part of the pontine nuclei (**Figure 7D,E; Supplementary Figure 3G,H**). By contrast, tracer injections in *Nex-cKO* brains, at locations corresponding to control S1 representations of the whiskers or upper limb, or into the S1/M1 (sensorimotor) lower limb representation, gave rise to abnormal distribution of corticopontine fibres (**Figure 7D-G**). Specifically, the corticopontine projections from medial parts of parietal cortex (corresponding to the S1 forelimb and hindlimb representations in control mice) were shifted towards more rostral locations in *Nex-cKO* experiments (**Figure 7D-F**, red points), resembling the distributions observed after tracer injections into motor areas in the control experiments (**Figure 6D-F**).

Notably, tracer injections placed in the anterolateral parietal cortex in *Nex-cKO* mice, in regions normally representing sensory surfaces of the head, gave rise to labelled axons distributed in the central part of the pontine nuclei, with more subtle difference to the matching control experiments (**Figure 7G**). In two cases, projections from regions corresponding to the S1 head representation in control mice were distinctly shifted towards medial relative to control experiments (red dots in **Figure 7G, and I**), attaining a distribution resembling the corticopontine projections from head representations in M1 cortex, located significantly more anteriorly in the cortex (grey dots in **Figure 7I**). Interestingly, two tracer injections in similar locations, corresponding to the border between the nose and whisker representations, gave rise to labelling located in central and rostral parts of the pontine nuclei (**Figure 7F,G**, red dots, see also **Figure 7C’**) with predominantly rostrally-shifted fibres in one case (**Figure 7F,C’; Supplementary Figure 5J**), and centrally-located and medially-shifted fibres in the other case (**Figure 7G; Supplementary Figure 5K**). Despite the differences in density distributions, both cases represent a distinct shift of fibre distributions towards regions normally receiving motor projections. Taken together, our observations indicate that corticopontine projections from anterolateral cortex in *Nex-cKO* mice also display abnormal topographical distributions resembling the normal projections from homologous representations in the more anterior and medially located primary motor cortex of control mice. Finally, one tracer injection placed in the most anterolateral part of S1 in a *Nex-cKO* mouse, at a location corresponding to the perioral surface representations in controls, yielded corticopontine labelling which was highly similar to that of a control experiment (**Figure 7H**).

Our findings thus show that corticopontine projections from frontal (motor) areas and the most anterolaterally located parts of the parietal (S1) cortex are topographically similar in *Nex-cKO* brains and controls, whereas corticopontine projections from most parts of parietal cortex, where the head, whisker, upper limb, and lower limb representations of S1 are located in control mice, are abnormally shifted towards rostral and medial regions of the pontine nuclei, that normally receive projections from cortical motor areas (schematically summarized in Fig. 7J,K). This is in overall agreement with the spatial and temporal control of Nr2f1 in area mapping. Indeed, the absence of changes in corticopontine projections from the frontal (motor) cortex might be due to low Nr2f1 expression in this area (spatial control), whereas lack of changes in the projections originating from the most anterolateral part of the parietal (S1) cortex, from which the earliest cortical projections to innervate the forming pontine nuclei originate, might be explained by the late Nr2f1 genetic inactivation occurring after the earliest layer V neurons have been produced (temporal control).

## Discussion

Our present study questions whether and how spatial-temporal cortical expression gradients are involved in the establishment of normal topographical organization of corticopontine projections. By combining genetically-modified mice and public mouse brain connectivity data with tract-tracing techniques and digital brain atlas tools, we have provided novel evidence of an intrinsic molecular control of layer V cortical neurons during the establishment of topographical organization of corticopontine projections. Abnormal areal organization in the neocortex induced by *Nr2f1* inactivation is reflected in altered corticopontine projections, as well as impaired structural integrity of the CST. While loss of *Nr2f1* from the early progenitor cell pool leads to increased and abnormal corticopontine innervation at the expense of corticospinal projections, only late postmitotic *Nr2f1* inactivation reveals altered topographic pontine mapping from medially located parts of somatosensory cortex controlling whiskerand limb representations. No shifts in projections from the earliest generated anterolateral cortical areas were observed in these mice, in line with a spatial and temporal control of Nr2f1 expression, respectively. Overall, our data show that proper area mapping of the neocortical primordium is a pre-requisite for preserving the cortical spatial and temporal segregation within the pontine nuclei, and thus correct corticopontine topographic organization.

### Spatial accuracy of topographical data compared across experiments

To ensure accurate 3D data in wild-type and genetically-modified mice, we relied on spatial alignment of serial microscopic section images to a common reference atlas achieved through non-linear image registration method (Puchades et al., 2019). The use of non-linear registration compensated for minor shape differences among brains and allowed comparison of distribution patterns among spatially relevant data. The focus on the location rather than the amount of signal expression/axonal labelling also compensated for the variation in signal expression intensity and size of tracer injections among cases. By representing signal expression and axonal labelling as 3D point clouds, it became possible to directly explore and compare location and distribution patterns in 3D in different combinations of data sets. For the additional benchmark data extracted from the *Allen Mouse Brain Connectivity Atlas, we* used the same sagittal image orientation as in our microscopic data, to facilitate comparison of microscopic images in addition to the 3D comparisons. The relevance and accuracy of the approach was confirmed by demonstrating that similarly located cortical tracer injections in control animals gave rise to similarly distributed labelling patterns in the pontine nuclei. Given the distinct patterns of topographical organization of corticopontine projections, the interpretation and comparison of spatial distribution patterns and variability in our tract tracing experiments critically depended on the analysis of tracer injection locations, and a 3D understanding of the topographical mapping of the cortical surface onto the pontine nuclei, derived from our analyses of control experiments and earlier studies of the rat corticopontine system (Leergaard and Bjaalie, 2007).

### Mitotic versus postmitotic Nr2f1 functions in layer V corticofugal projections

Our previous work showed overall areal organization impairments in cortical *Nr2f1* mutant brains, independent of *Nr2f1* inactivation in progenitors or postmitotic neurons. Here, we report for the first time that *Nr2f1* drives corticopontine connectivity differently in progenitors *versus* postmitotic neurons. While *Nr2f1* expressed by progenitor cells modulates the ratio between corticopontine and corticospinal axonal projections, similarly to what happens in *C. elegans* with the ortholog *UNC-55* (Petersen et al., 2011; Zhou and Walthall, 1998), postmitotic *Nr2f1* expression specifically acts on somatosensory topographic organization of corticopontine neurons (**Figures 4, 5**). This suggests that early *Nr2f1* expression in progenitor cells is mainly required in the initial axonal pathfinding of layer V subtypes, while later postmitotic expression is more implicated in the later refinement of corticopontine topographical organization. Since YFP+ cells follow the physiological distribution of subcortical layer V projections (Porrero et al., 2010), Thy1-driven YFP fluorescence will be induced in cortical subpopulations related to the specificity of their axonal target region; this tight relationship which will not be altered in the conditional *KO* models. Accordingly, a higher number of layer V neurons in parietal *Emx1-cKO* cortices leads to disorganized corticopontine innervation, in accordance with increased Lmo4 expression, known to drive layer V neurons *versus* the pontine nuclei (Cederquist et al., 2013; Harb et al., 2016). Differently, *Nex-cKO* parietal axonal projections reach pontine targets normally innervated by motor-derived cortical areas. Finally, increased number of YFP+ cells in visual and auditory areas in the occipital cortex, corresponds with an augmented innervation in dorsolateral regions of pontine nuclei known to receive projections from the occipital cortex. Together, these data indicate a dual role for Nr2f1 in layer V corticofugal connectivity, an early role in subtype specification (corticopontine vs corticospinal), and a later role in topographical mapping.

### Revising the chrono-architectonic hypothesis of cortico-pontine circuit development

Previous data in developing rats have shown that pontine neurons settle in the forming pontine nuclei in a shell-like fashion according to their birthdate, with early born neurons forming the central core of the pontine nuclei, and later born neurons consecutively settling around the earlier born neurons forming concentric rings (Altman and Bayer, 1987). At early postnatal stages, corticopontine axons are chemotropically attracted as collateral branches from corticospinal axons (Heffner et al., 1990; O’Leary and Terashima, 1988), innervating the pontine nuclei in a topographic inside-out pattern (Leergaard et al., 1995). Neurons in the frontal (motor) cortex project rostrally and medially in the pontine nuclei, neurons in the parietal (somatosensory) cortex project to central and caudal parts, neurons in the temporal (auditory) cortex to central and lateral regions, and neurons in the occipital (visual) cortex to lateral and rostral parts of the pontine nuclei (Leergaard and Bjaalie, 2007). This concentric organization of corticopontine projections suggests that the birthdate of pontine neurons and their inside-out genesis is linked to the spatial organization of cortical inputs. However, intrinsic differences in pontine neurons born at different times might also have an instructive role for corticopontine innervation. A recent study in mice showed that postmitotic expression of the HOX gene *Hoxa5* guides pontine neurons to settle caudally within the pontine nuclei, where they are targeted by projections from limb representations in the somatosensory cortex (Maheshwari et al., 2020). Moreover, ectopic *Hoxa5* expression in pontine neurons is sufficient to attract cortical somatosensory inputs, regardless of their spatial position, showing that pontine neurons can play an instructive and attractive role in topographic input connectivity of corticopontine neurons (Maheshwari et al., 2020).

Nevertheless, maturational gradients in the pontine nuclei cannot fully explain the complexity of the fine-grained somatotopic topographic connectivity pattern between cortical input and pontine neuron targets. Since the establishment of topographic maps requires multiple processes and structures, it is conceivable that the position and specific intrinsic molecular programs of both presynaptic afferents and postsynaptic target neurons contribute to this complex corticopontine connectivity map. Indeed, our data show that without affecting the development and maturation of pontine neurons, corticopontine Nr2f1-deficient layer V axons originating from the parietal S1 cortex will abnormally target the pontine region normally deputed to corticopontine motor axons. By contrast, Nr2f1-deficient axons originating from the frontal and medial cortex will innervate the expected pontine region allocated to motor axons. This strongly suggests that during the establishment of corticopontine topography, both structures, the neocortex and the pons need to be properly pre-patterned by factors involved in spatial and temporal control of neurogenesis, such as *Nr2f1* for the cortex, and *Hoxa5* for the pontine nuclei (model in **Figure 8**).

**Figure 8.**
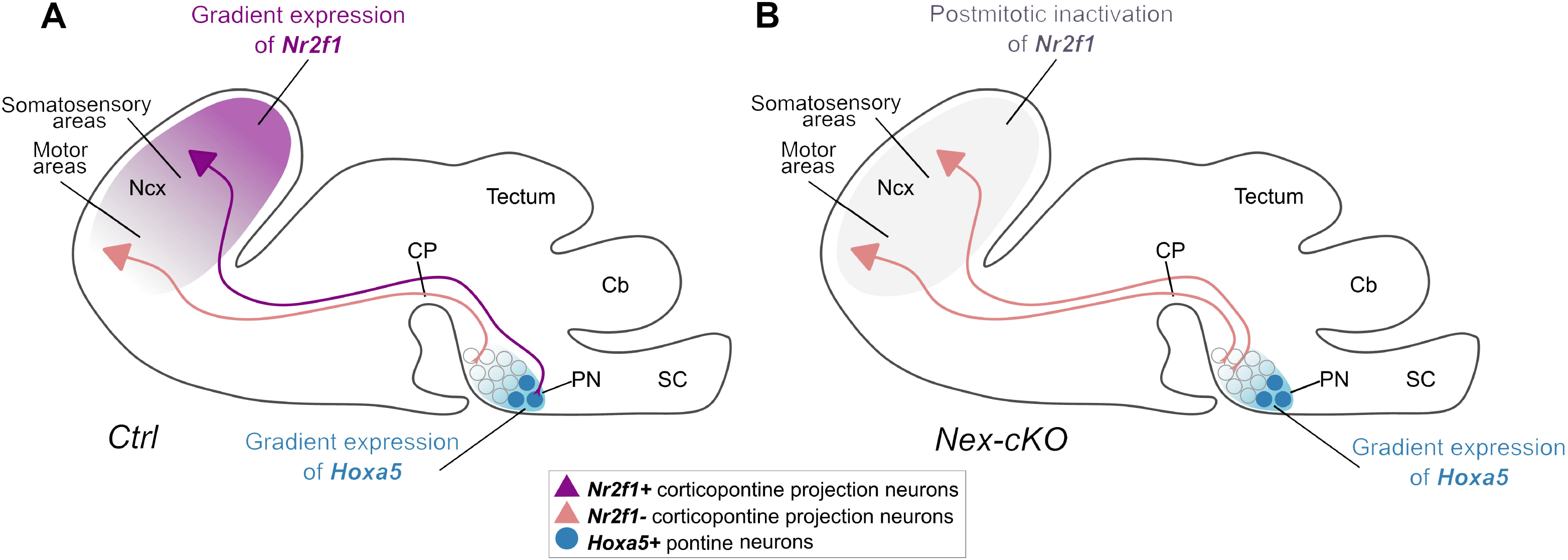
Model of corticopontine topography establishment and changes upon Nr2f1 cortical abolishment. (**A**) Proposed schematic model of how the neocortex (Ncx) and the pontine nuclei (PN) might interact during development in driving corticopontine topography. Both structures need to be pre-patterned by gradient expression of transcription factors. While postmitotic pontine graded expression of Hoxa5 will impart an anterior to posterior identity to pontine neurons (graded blue circles) (Maheshwari et al., 2020), postmitotic cortical gradient expression of Nr2f1 (purple gradient in Ncx) will intrinsically instruct corticopontine neurons to innervate their topographically proper targets (this study). (**B**) In the absence of postmitotic Nr2f1 gradient expression in the neocortex, but maintenance of Hoxa5 expression in the PN, axons from motor and somatosensory cortex will both innervate rostrally-located clusters within the PN, since somatosensory corticopontine projections in Nex-cKO mice are changed to resemble motor-like projections. Abbreviations: Cb, cerebellum; CP, cerebral peduncle; SC, spinal cord.

### Conclusion and outlook

By showing that gradient cortical expressions of transcription factor Nr2f1 is directly involved in the establishment of corticopontine topographic mapping, we provide new insights into the development of neural systems. However, other factors regulating the size and location of cortical areas also are likely also implicated. We conclude that distinct molecular mechanisms in the source (cerebral cortex) and target (pontine nuclei) regions must be coordinated during the establishment of corticopontine topography. Identifying the molecular pathways within the cortex and pontine nuclei, as well as the mechanisms and molecules governing their interaction remains an open question for further studies.

### Limitation of the study

This study shows a limited number of viral injections per cortical area that was partially compensated by adapting mouse tract-tracing data from the Allen Brain Institute to our analysis. Even through both groups of samples showed a highly consistent pattern of topographical organization, the two types of data resulted from different experimental conditions. Because of some variability of the fluorescent signal among samples, we chose to focus on 3D spatial distribution patterns that turned to be consistent across all control experiments. We found the *Emx1-cKO* mouse model less suitable for further topographical analysis due to the disorganized and chaotic innervation pattern observed in the *Thy1-eYFP-H* reporter background. Although more sophisticated methods are needed to pinpoint the cellular and molecular mechanisms involved in the establishment of corticofugal topography, our study represents a useful starting point and resource for further studies of the development of corticopontine and corticospinal projections in mice.

## Materials and Methods

### Topographical map of corticopontine projections from somatosensory and motor areas

To establish a 3D benchmark map of corticopontine projections from somatosensory and motor areas in in adult wild type mice, we utilized a selection of public experimental tract-tracing data available from the Allen Institute mouse brain connectivity atlas (http://connectivity.brain-map.org/). We selected 11 experiments in which the anterograde tracer EGFP was injected in the right primary/secondary motor cortex (n =6) or primary somatosensory cortex (n = 5) of wild type C57BL/6J mice (**Supplementary Table 1**). Serial two-photon fluorescence images were interactively inspected using the Projection High Resolution Image viewer of the Allen Institute, and from each case, 5 sagittal oriented images of the right pontine nuclei spaced at ~100 μm were captured by screen shot from the largest 3D multiplane thumbnail viewer. The resolution of the captured images was up-sampled three times original size before their spatial alignment to the CCFv3 was optimized using the tools QuickNII (Puchades et al., 2019) and VisuAlign (RRID: SCR_017978), as described below. These images were used to create 3D representations of the axonal labelling in the pontine nuclei (**Figure 2**; see below).

### Animals

All mice used were bred in a C57BL/6J background. Male and female animals at any stage of development were used. All experiments were conducted in accordance with the French Animal Welfare Act and European guidelines for the use of experimental animals, using protocols approved by the French Ministry of Education, Research and Innovation and the local ethics committee (CIEPAL NCE/2019-548, Nice) under authorization #15 349 and #15 350. *Nr2f1/C0UP-TFI^fl/fl^* mice were crossed with *Emx1-Cre-recombinase* mice to inactivate *Nr2f1/COUP-TFI* exclusively in cortical progenitors and their progeny (Armentano et al., 2007) or with *Nex-Cre-recombinase* mice to abolish *Nr2f1/COUP-TFI* expression from postmitotic neurons (Alfano et al., 2014). Littermate *Nr2f1/COUP-TFI^fl/fl^* mice without the presence of the *Cre-recombinase* gene (*Cre-negatives*) were considered controls (**Supplementary Table 2**). For postnatal (P)21 and adult topographic map analysis, *Emx1-cKO* and *Nex-cKO* animals were further crossed with *Thy1-eYFP-H* mice to specifically label layer V projection neurons, as previously reported (Harb et al., 2016; Porrero et al., 2010). Mice were genotyped as previously described (Alfano et al., 2014; Armentano et al., 2007; Harb et al., 2016). Control and mutant littermates were genotyped as *Nr2f1^fl/fl^:Thy1-eYFP-H^T/+^* and *Nr2f1^fl/fl^:Emx1-Cre:Thy1-eYFP-H^T/+^* or *Nr2f1^fl/fl^:Nex-Cre:Thy1-eYFP-H^T/+^*, respectively. For simplicity, mutant mice are named *Emx1-cKO* and *Nex-cKO* throughout the text. Midday of the day of the observed vaginal plug was considered as embryonic day 0.5 (E0.5).

### Anterograde tracing of corticospinal axons in early postnatal mice

P4-P5 animals were anesthetized on ice for 5 min and kept on ice during the whole procedure. Viral particles were produced from the AAV9-CAGtdTomato plasmid by the Alexis Bemelmans (CEA, France) Company, and diluted 1:50 in TE-Buffer (Qiagen, #1018499) to a final concentration of 1.75e12 vg/ml (kindly donated by I. Dusart, Pierre and Marie Curie University, Paris, France). Approximately 0.5/1ul was injected unilaterally in different rostral-caudal and medio-lateral brain locations of control and *Nex-cKO* pups, as previously described in (Gu et al. 2017).

### Microscopic imaging

Mosaic microscopic images were acquired using an Axio Imager M2 epifluorescence microscope (Carl Zeiss Microscopy GmbH, Jena, Germany) equipped with a halogen lamp, a MCU 2008 motorized stage, and an EC Plan-Neofluar 10x/0.30 and an AxioCam MRm camera. ZEN blue software was used for imaging and automatic stitching. Images were exported in TIFF format and serially ordered from lateral to medial, rotated and if needed, mirrored to consistent anatomical orientation using Adobe Photoshop CS6 (RRID: SCR_014199), before being converted to PNG format and resized to 60% of original size using ImageJ (RRID: SCR_003070) with bilinear interpolation. The resized serial images were loaded into Adobe Photoshop as a stack, spatially aligned using the ventral surfaces of the pons and cerebral peduncle as landmarks, cropped and exported as individual PNG files. For comparative analyses of topographical organization (see below), variations in YFP signal expression intensity within and between groups were normalized by adjusting the brightness and contrast of images to equal levels using a custom-made histogram matching script available for ImageJ (National Institutes of Health; https://imagej.nih.gov/). One selected, representative case (Experiment 5, Cre-negative, nr: 14250, **Supplementary Table 2**) was used as reference.

### Spatial alignment to common 3D reference atlas

Serial sectional images were spatially registered to the *Allen Mouse Common Coordinate Framework*, version 3, 2017 edition of the delineations (CCFv3, (Wang et al., 2020) using the QuickNII software tool (RRID:SCR_016854; (Puchades et al., 2019). Multiple anatomical landmarks (hippocampus, caudate-putamen, inferior and superior colliculus, and external surface of the neocortex) were used to determine the medio-lateral position and orientation of the sagittal section images. For each section image, custom atlas diagrams were aligned to anatomical landmarks in the experimental images using affine transformations, with emphasis on matching the ventral surface of the pons and white matter tracts close to the pontine nuclei and exported as PNG images. As a secondary step, to further optimize registration, the custom atlas images were non-linearly transformed using the software tool VisuAlign v0.8 (RRID:SCR_017978), with particular focus on fitting the template to the outer brain surface, subcortical white matter, and the outer boundaries of the pontine nuclei. To co-display images and the spatially registered custom atlas images, we used the software tool LocaliZoom, which is embedded in the Navigator3 image management system (bit.ly/navigator3), developed and hosted by the Neural Systems Laboratory at the University of Oslo, Norway.

### Cortical distribution analysis in *Emx1-cKO* and *Nex-cKO* mutants

Serial section images from *Nex-cKO* and *Emx1-cKO* mutants co-registered to *the Allen Mouse Brain Connectivity Atlas* were analysed using an ImageJ macro allowing automatic counting of spots by area of interest in the brain. The spots are considered as specific staining using a threshold based on intensity and shape of the elements and the composite RGB segmentation atlas as a mask for the region of interest. The macro pre-processes an atlas plate to delineate the regions of interest (ROI), based upon their unique colour-code and corrects the corresponding raw image by subtracting the background generating binary images of the signal and combining it to the ROI maps. Finally, all objects within circularity range of 0.5-1 were counted per ROI using the *find maxima* tool. The process, reiterated for each atlas plate-raw image combination, produced a summary table containing the quantification of particles per area per section, shown as graphs in Figure 3.

### Corticospinal tract morphometric analysis in *Emx1-cKO* and *Nex-cKO* mutants

Serial section images from *Nex-cKO* and *Emx1-cKO* mutants were analysed using the *Fiji-ImageJ Software* tool (Schindelin et al., 2015) to determine the total dorsoventral width of the bundle expressing fluorescent signal in the descending fibre tract in different positions: rostrally and caudally to the pontine nuclei, and 250 μm and 500 μm caudal to the nuclei. The width of separate fibre fascicles was also measured 250 μm and 500 μm from the terminal edge of the pontine nuclei (**Figure 4A’**).

### Analysis of tracer injection sites

Serial section images of cortical tracer injections in *Nex-cKO* brains (**Supplementary Table 3**) and experiments taken from the *Allen Mouse Brain Connectivity Atlas*, were spatially aligned using QuickNII and VisuAlign, as described above. The centre positions of the injection sites were annotated as a point-coordinate using LocaliZoom, co-displayed with the CCFv3 atlas in the 3D viewer tool MeshView (RRID:SCR_017222). These visualizations were used to select spatially corresponding injection site locations for analyses of spatial distribution of corticopontine projections.

### Histology, immunohistochemistry and *in situ* hybridization

At P21 and adulthood, animals were anesthetized by intraperitoneal injection of a mixture of Tiletamine-Zolazepam-Xylazine-Buprenorphine and intracardially perfused with PB Saline (PBS) followed by 4% paraformaldehyde (PFA) in PBS. Volumes were 15, 20 and 30 ml, respectively. Brains were removed from the skull and postfixed for 4 h at 4°C in 4% PFA, before being vibratome-sectioned in 100μm (adult samples) or 150μm (P21 samples) thick sagittal sections. All sections were incubated overnight at 4°C in a solution of 0.5% Triton X-100, 3% BSA, 10% goat serum in PBS, for permeabilization and reduction of non-specific binding of antibodies. For immunofluorescence (IF), sections were incubated for 2 days at 4°C with primary antibodies in a solution of 0.5% Triton X-100, 10% goat serum in PBS, and then overnight at 4°C with relative secondary antibodies and HOECHST diluted in PBS. For the complete list of primary and secondary antibodies, see **Supplementary Table 5**. Sections were washed several times in PBS, then transferred on Superfrost plus slides (ThermoScientific), covered and dried for 30 min to 1 h, and finally mounted with the Mowiol (Sigma-Aldrich) mounting medium.

### Semi-quantitative recording and 3D visualization of spatial distribution patterns

To investigate and compare the 3D distributions of YFP signal expression or anterograde axonal labelling within the pontine nuclei, we used the annotation functionality in the LocaliZoom tool to semi-quantitatively record YFP signal expression or labelled axons in all sections through the pontine nuclei as point coordinates (specified in the coordinate system of the reference atlas, CCFv3), reflecting the overall density of signal/labelling observed in the images (**Figure 2C-D**). To compensate for the spacing between sections and allow inspection of point distributions perpendicularly to the section angle, the z-coordinate of each point was randomly displaced within the thickness of the gap between sections using a custom Python script. The point-coordinates were co-displayed in the MeshView 3D viewer tool (**Figures 2, 5, 6, 7**).

## Acknowledgements

We thank Isabelle Dusart in Paris for providing us with AAV9-CAG-tdTomato viral particles produced by Alexis Bemelmans (CEA, France), Nicolaas Groeneboom and Gergely Csucs for expert technical assistance, Baptiste Monterroso and Sameh Ben Aicha of the iBV PRISM microscopy platform for the set-up and optimization of automatic image analysis tools used in this work, and Sharon Yates, and Maja A. Puchades for useful discussions.

## Data and Code Availability

All microscopic images generated in this project will be shared via the EBRAINS research infrastructure (https://search.kg.ebrains.eu/) as high-resolution TIFF images under a CC-BY license, with a unique DOI for each dataset. All tract tracing image data downloaded from the Allen Institute mouse brain connectivity atlas (Supplementary Table 1) are publicly available from http://connectivity.brain-map.org/. The custom Python script (spread.py) used to randomly displace data points within the thickness of a section is available upon request (to author JGB). The QuickNII (v2.2, RRID: SCR_016854; Puchades et al., 2019) LocaliZoom and Meshview (RRID: SCR_0170222) tools are available from EBRAINS (https://search.kg.ebrains.eu/).

