## Supplementary Figures and Tables for "Topography of corticopontine projections is controlled by postmitotic expression of the area-mapping gene Nr2f1"

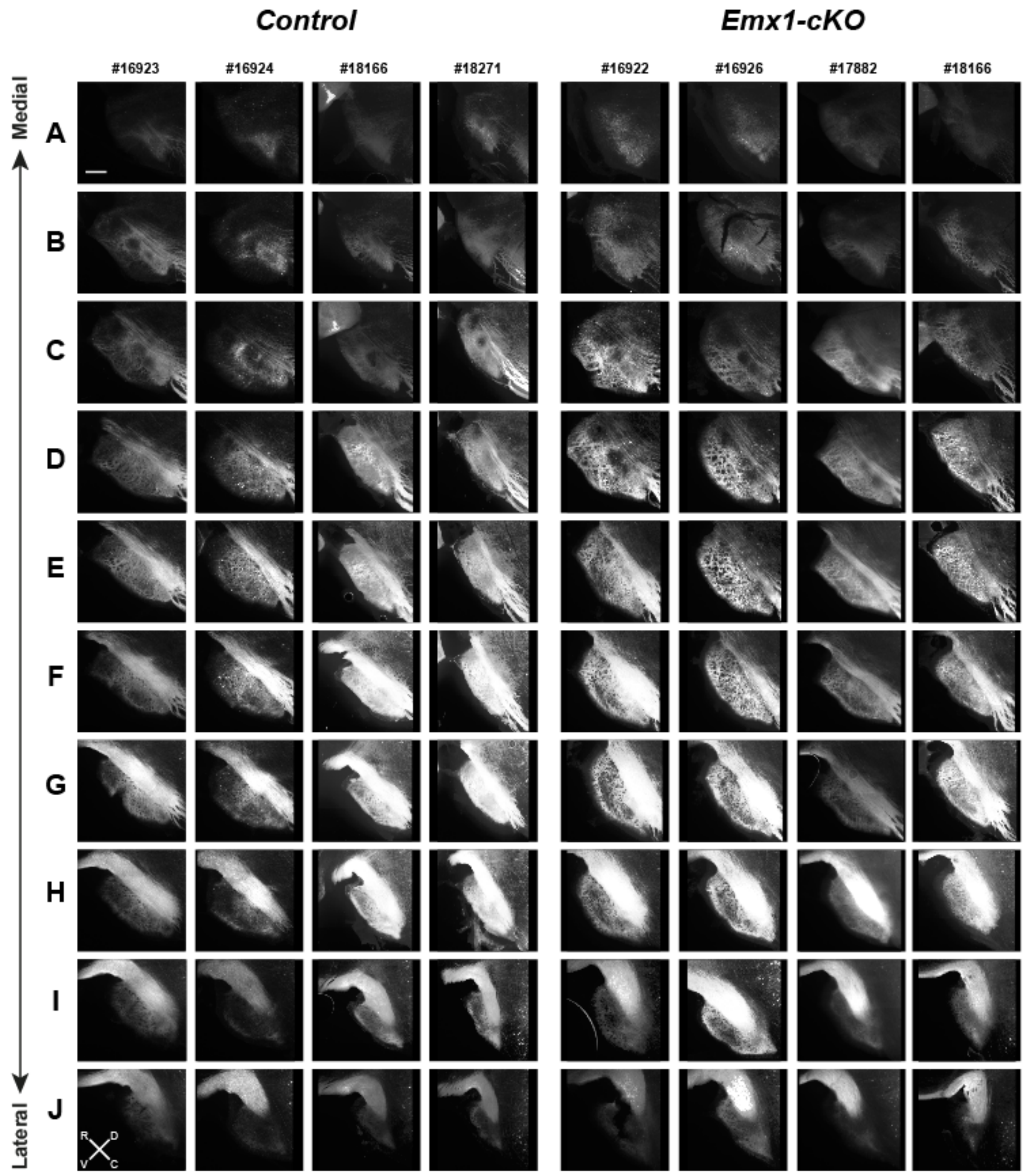

**Supplementary Figure 1. Overview of YFP-expression in *Emx1-cKO* mice and controls**

*Fluorescence microscopy images of the pontine nuclei in sagittal sections from 4 control- and 4 Emx1-cKO mice. Columns show images from one animal, with sections from corresponding levels from medial to lateral are sorted from top to bottom (rows A-J). The intensity levels of the images have been normalized. Signal expression in Emx1-cKO mice is more widespread and more diffusely distributed in the pontine nuclei, relative to controls. Scale bar, 200 µm.*

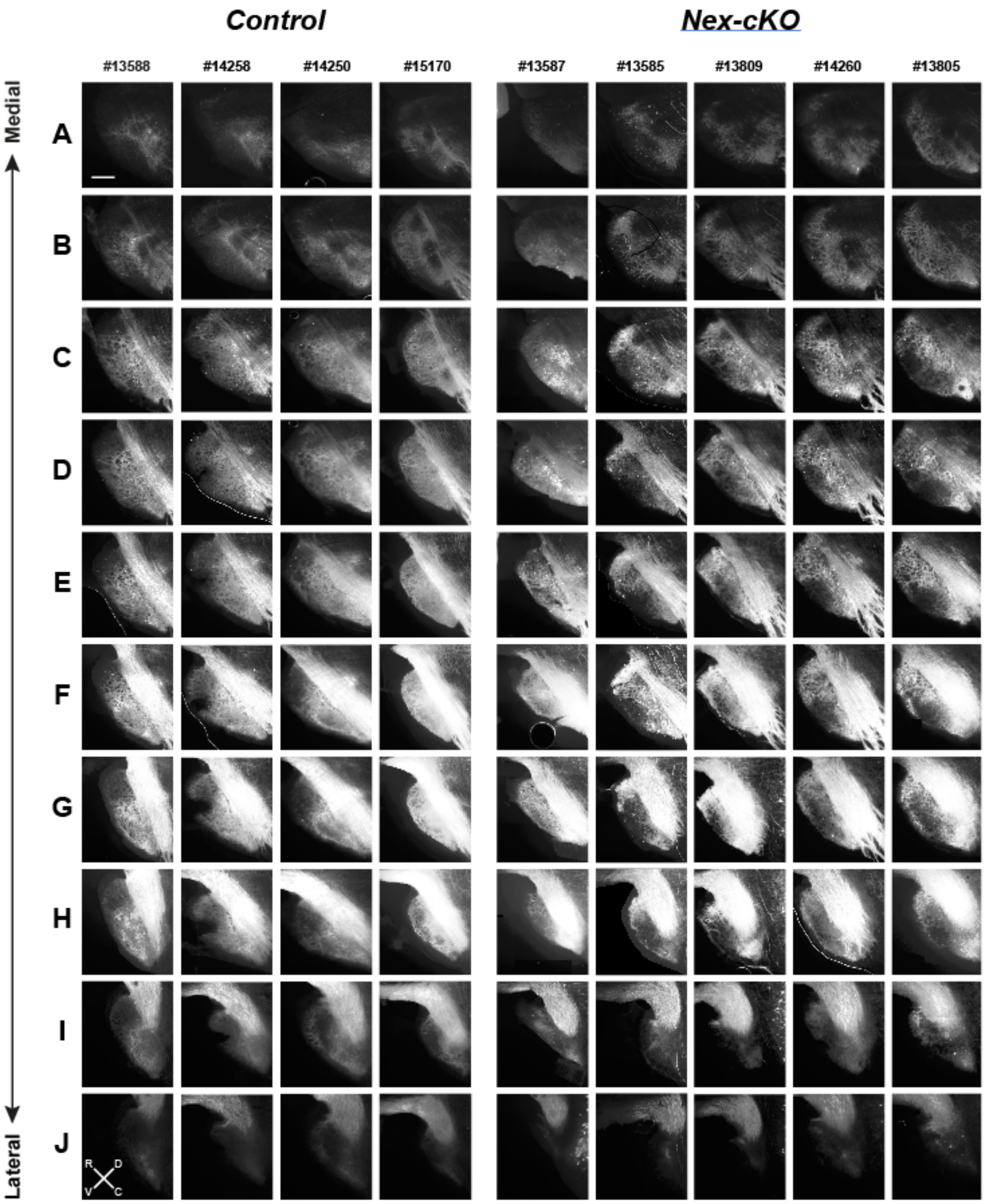

**Supplementary Figure 2. Overview of YFP-expression in *Nex-cKO* mice and controls**

*Fluorescence microscopy images of the pontine nuclei in sagittal sections from 4 control- and 4 Nex-cKO mice. Columns show images from one animal, with sections from corresponding levels from medial to lateral are sorted from top to bottom (rows A-J). The intensity levels of the images have been normalized. Signal expression in Nex-cKO mice is more clearly reduced or absent in the central core region of the pontine nuclei, relative to controls. Scale bar, 200 µm.*

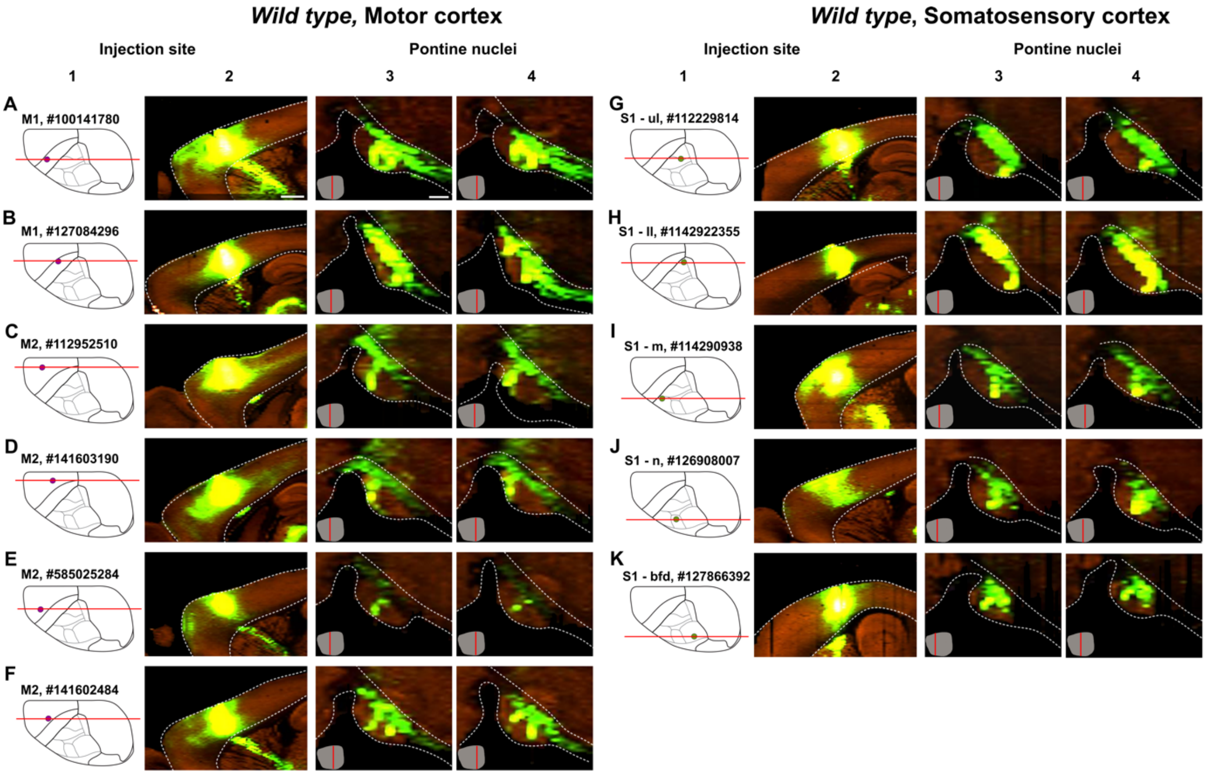

**Supplementary Figure 3. Overview of tract-tracing experiments in wild type mice**

*(**A-K**) Sagittally-oriented fluorescence microscopy images showing all wild-type C57BL/6J mouse tract-tracing data captured from the online image viewer of the Allen Mouse Brain Connectivity Atlas. **Column 1** shows dorsal view diagrams of the cerebral indicating the position of the tracer injection site as a red dot, and a red line indicating the location of the sagittal images in **column 2** that show injection site centres. Letters and numbers indicate injected cortical area and ID numbers. **Columns 3 and 4** show fluorescence microscopy images of sagittal sections through the pontine nuclei, showing representative corticopontine labelling at two mediolateral levels as indicated with red lines on the inset ventral view diagram of the pontine nuclei. Abbreviations, bfd, barrel field; M1, primary motor cortex; M2, secondary motor cortex; ll, lower limb; m, mouth; n, nose; ul, upper limb. Scale bars, 1mm (column 2), 200 µm (columns 3, 4).*

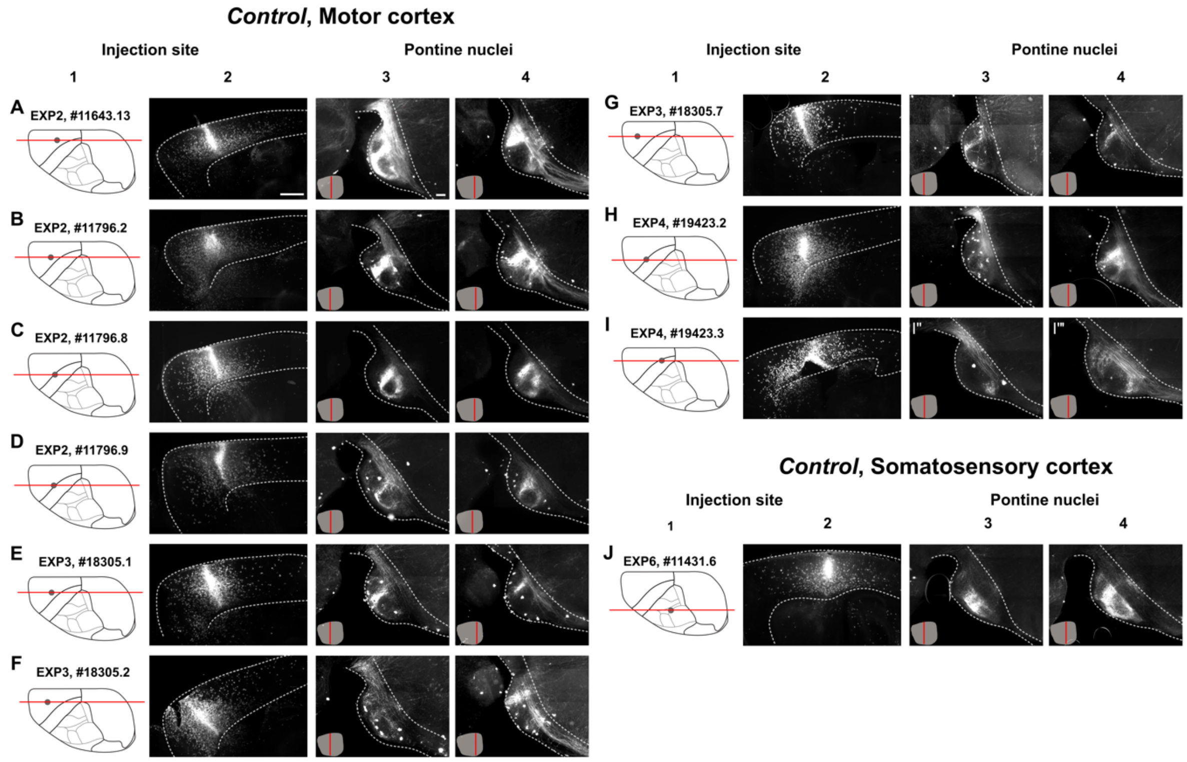

**Supplementary Figure 4. Overview of tract-tracing experiments in control mice**

*(**A-J**) Sagittally-oriented fluorescence microscopy images showing all tract-tracing data conducted in control animals. **Column 1** shows dorsal view diagrams of the cerebral indicating the position of the tracer injection site as a red dot, and a red line indicating the location of the sagittal images in **column 2** that show injection site centres. Letters and numbers indicate injected cortical area and ID numbers. **Columns 3 and 4** show fluorescence microscopy images of sagittal sections through the pontine nuclei, showing representative corticopontine labelling at two mediolateral levels as indicated with red lines on the inset ventral view diagram of the pontine nuclei. Scale bars, 1mm (column 2), 200 µm (columns 3, 4).*

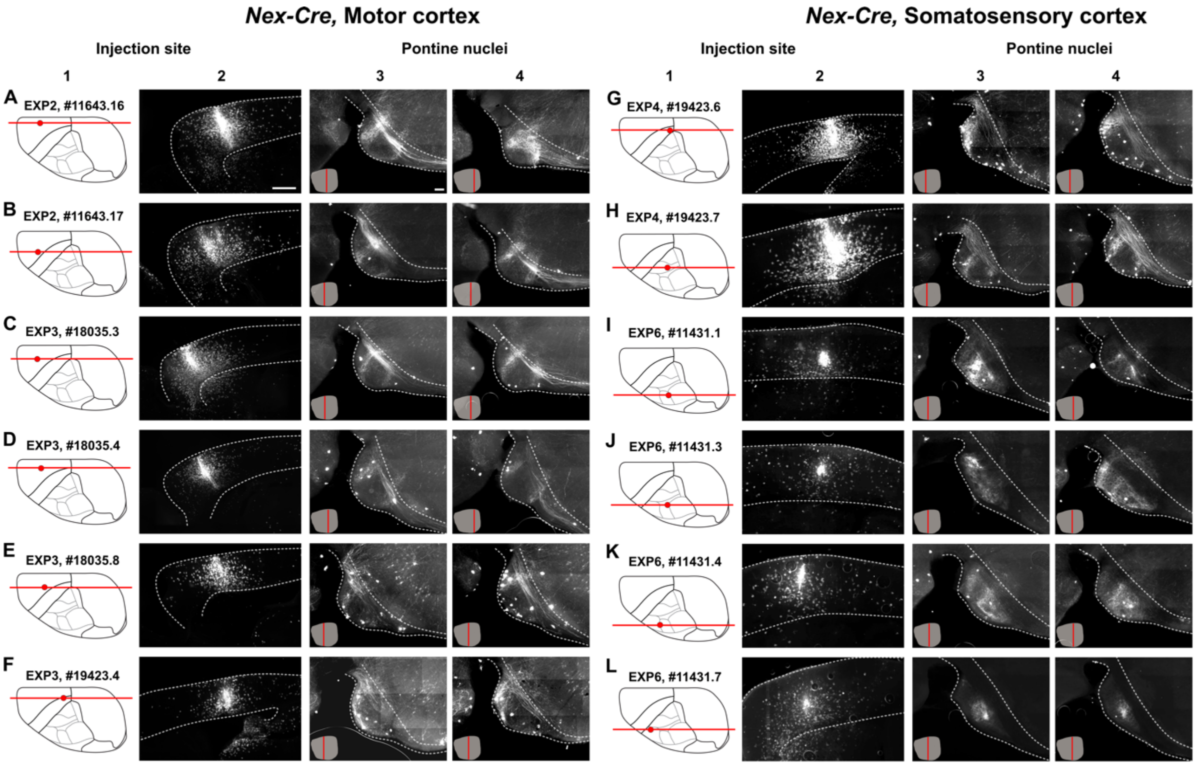

**Supplementary Figure 5. Overview of tract-tracing experiments in *Nex-cKO* mice**

*(**A-L**) Sagittally-oriented fluorescence microscopy images showing all tract-tracing data conducted in Nex-cKO animals. **Column 1** shows dorsal view diagrams of the cerebral indicating the position of the tracer injection site as a red dot, and a red line indicating the location of the sagittal images in **column 2** that show injection site centres. Letters and numbers indicate injected cortical area and ID numbers. **Columns 3 and 4** show fluorescence microscopy images of sagittal sections through the pontine nuclei, showing representative corticopontine labelling at two mediolateral levels as indicated with red lines on the inset ventral view diagram of the pontine nuclei. Scale bars, 1mm (column 2), 200 µm (columns 3, 4).*

### SUPPLEMENTARY TABLES

#### *Supplementary Table 1. Overview of wild-type experiments from the Allen Mouse Brain Connectivity database*

| **Allen Mouse Brain Connectivity database** | | | | | |
|---|---|---|---|---|---|
| **Experiment number #** | **Sex** | **Age (±2)** | **Genotype** | **Injection site** | **Shown in** |
| 100141780 | Male | P56 | C57BL/6J | Primary motor cortex | Fig. 2, 6A-F |
| 114290938 | male | P56 | C57BL/6J | Primary somatosensory cortex, mouth region | Fig. 2A-P, 7A and 7H |
| 112229814 | male | P56 | C57BL/6J | Primary somatosensory cortex, upper limb region | Fig. 2, 7B and 7F |
| 112952510 | male | P56 | C57BL/6J | Secondary motor cortex | Fig. 2 |
| 114292355 | male | P56 | C57BL/6J | Primary somatosensory cortex, lower limb region | Fig. 2, 7A and 7D |
| 126908007 | male | P56 | C57BL/6J | Primary somatosensory cortex, nose region | Fig. 2, 7A and 7G |
| 127084296 | male | P56 | C57BL/6J | Secondary motor cortex | Fig. 2 |
| 127866392 | male | P56 | C57BL/6J | Primary somatosensory cortex, barrel field region | Fig. 2 |
| 141602484 | male | P56 | C57BL/6J | Secondary motor cortex | Fig. 2 |
| 141603190 | male | P56 | C57BL/6J | Secondary motor cortex | Fig. 2 |
| 585025284 | male | P56 | C57BL/6J | Secondary motor cortex | Fig. 2 |

***Supplementary Table 2. Overview of Emx-cKO and Nex-cKO mice***

| **Adult** | | | | | |
|---|---|---|---|---|---|
| **Exp. #** | **Animal #** | **Sex** | **Age** | **Genotype** | **Shown in** |
| 1 | 13588 | female | P33 | *Thy1-eYFP*$^{T/+}$*; Nr2f1*$^{fl/fl}$ | Fig. 3, 4, 5 and Suppl. Fig.1, 2 |
| 1 | 13587 | female | P33 | *Thy1-eYFP*$^{T/+}$*; Nr2f1*$^{fl/fl}$*; Nex-Cre* | Fig. 3, 4, 5 and Suppl. Fig.1, 2 |
| 2 | 13585 | male | P62 | *Thy1-eYFP*$^{T/+}$*; Nr2f1*$^{fl/fl}$*; Nex-Cre* | Fig. 3, 4, 5 and Suppl. Fig.1, 2 |
| 3 | 13809 | male | P57 | *Thy1-eYFP*$^{T/+}$*; Nr2f1*$^{fl/fl}$*; Nex-Cre* | Fig. 3, 4, 5 and Suppl. Fig.1, 2 |
| 4 | 14258 | male | P57 | *Thy1-eYFP*$^{T/+}$*; Nr2f1*$^{fl/fl}$ | Fig. 3, 4, 5 and Suppl. Fig.1, 2 |
| 4 | 14260 | male | P57 | *Thy1-eYFP*$^{T/+}$*; Nr2f1*$^{fl/fl}$*; Nex-Cre* | Fig. 3, 4, 5 and Suppl. Fig.1, 2 |
| 5 | 13805 | male | P55 | *Thy1-eYFP*$^{T/+}$*; Nr2f1*$^{fl/fl}$*; Nex-Cre* | Fig. 3, 4, 5 and Suppl. Fig.1, 2 |
| 5 | 14250 | male | P72 | *Thy1-eYFP*$^{T/+}$*; Nr2f1*$^{fl/fl}$ | Fig. 3, 4, 5 and Suppl. Fig.1, 2 |
| 5 | 15170 | male | P75 | *Thy1-eYFP*$^{T/+}$*; Nr2f1*$^{fl/fl}$ | Fig. 3, 4, 5 and Suppl. Fig.1, 2 |
| 6 | 16922 | male | P76 | *Thy1-eYFP*$^{T/+}$*; Nr2f1*$^{fl/fl}$*; Nex-Cre* | Fig. 3, 4, 5 and Suppl. Fig.1, 2 |
| 6 | 16923 | male | P76 | *Thy1-eYFP*$^{T/+}$*; Nr2f1*$^{fl/fl}$ | Fig. 3, 4, 5 and Suppl. Fig.1, 2 |
| 6 | 16924 | male | P76 | *Thy1-eYFP*$^{T/+}$*; Nr2f1*$^{fl/fl}$ | Fig. 3, 4, 5 and Suppl. Fig.1, 2 |
| 6 | 16926 | male | P76 | *Thy1-eYFP*$^{T/+}$*; Nr2f1*$^{fl/fl}$*; Nex-Cre* | Fig. 3, 4, 5 and Suppl. Fig.1, 2 |
| 7 | 17882 | male | P72 | *Thy1-eYFP*$^{T/+}$*; Nr2f1*$^{fl/fl}$*; Nex-Cre* | Fig. 3, 4, 5 and Suppl. Fig.1, 2 |
| 8 | 18046 | female | P109 | *Thy1-eYFP*$^{T/+}$*; Nr2f1*$^{fl/fl}$*; Nex-Cre* | Fig. 3, 4, 5 and Suppl. Fig.1, 2 |
| 8 | 18166 | female | P98 | *Thy1-eYFP*$^{T/+}$*; Nr2f1*$^{fl/fl}$ | Fig. 3, 4, 5 and Suppl. Fig.1, 2 |
| 8 | 18271 | female | P87 | *Thy1-eYFP*$^{T/+}$*; Nr2f1*$^{fl/fl}$ | Fig. 3, 4, 5 and Suppl. Fig.1, 2 |
| 9 | 19606 | female | P86 | *Thy1-eYFP*$^{T/+}$*; Nr2f1*$^{fl/fl}$*; Nex-Cre* | Fig. 4 |
| 9 | 19607 | female | P86 | *Thy1-eYFP*$^{T/+}$*; Nr2f1*$^{fl/fl}$*; Nex-Cre* | Fig. 4 |

***Supplementary Table 3. Overview tract tracing experiments in Nex-cKO and Ctrl mice***

| P21 – unilateral CST tracing | | | |
|---|---|---|---|
| *Tracer injected in motor cortex* | | | |
| Experiment # | Animal # | Genotype | Shown in |
| 2 | 11643_13 | *Ctrl* | Fig. 6A and 7B |
| 2 | 11643_16 | *Nex-cKO* | Suppl. Fig. 3 |
| 2 | 11643_17 | *Nex-cKO* | Fig. 6A and 6F |
| 2 | 11796_2 | *Ctrl* | Suppl. Fig. 3 |
| 2 | 11796_8 | *Ctrl* | Suppl. Fig. 3 |
| 2 | 11796_9 | *Ctrl* | Suppl. Fig. 3 |
| 3 | 18035_1 | *Ctrl* | Fig. 6A and 6F |
| 3 | 18035_2 | *Ctrl* | Fig. 6A and 6E |
| 3 | 18035_7 | *Ctrl* | Suppl. Fig. 3 |
| 3 | 18035_3 | *Nex-cKO* | Fig. 6A and 6E |
| 3 | 18035_4 | *Nex-cKO* | Suppl. Fig. 3 |
| 3 | 18035_8 | *Nex-cKO* | Fig. 6A and 76D |
| 4 | 19423_2 | *Ctrl* | Suppl. Fig. 3 |
| 4 | 19423_3 | *Ctrl* | Suppl. Fig. 3 |
| 4 | 19423_4 | *Nex-cKO* | Suppl. Fig. 3 |
| 4 | 19423_5 | *Nex-cKO* | Suppl. Fig. 3 |
| *Tracer injected in somatosensory cortex* | | | |
| 4 | 19423_6 | *Nex-cKO* | Fig. 7A and 7D |
| 4 | 19423_7 | *Nex-cKO* | Fig. 7A and 7E |
| 6 | 11431_1 | *Nex-cKO* | Fig. 7A and 7G |
| 6 | 11431_3 | *Nex-cKO* | Fig. 7A and 7H |
| 6 | 11431_4 | *Nex-cKO* | Fig. 7A and 7F |
| 6 | 11431_6 | *Ctrl* | Fig. 7A and 7F |
| 6 | 11431_7 | *Nex-cKO* | Fig. 7A and 7H |

***Supplementary Table 4: Overview of quantitative results:***

*Highlighted in blue, comparisons that produced statistically significant P-values. P-values are calculated by 2way ANOVA test (**Figures 3F,G and 4E-F**), or ordinary one-way ANOVA test (**Figures 4G-I**).*

**Figure 3B"-D"– Cortical distribution of YFP-H positive cells (% values)**

| Area | Hypothesis | Mean 1 | Mean 2 | Mean diff. | 95,00% CI of diff | Summary | Adjusted P. |
|---|---|---|---|---|---|---|---|
| PFC | *Ctrl vs. Nex-cKO* | 4.833 | 1.95 | 2.883 | -3.655 to 9.421 | ns | 0.5437 |
| | *Ctrl vs. Emx1-cKO* | 4.833 | 3.15 | 1.683 | -4.855 to 8.221 | ns | 0.8112 |
| | *Nex-cKO vs. Emx1-cKO* | 1.95 | 3.15 | -1.2 | -8.362 to 5.962 | ns | 0.9150 |
| M | *Ctrl vs. Nex-cKO* | 46.87 | 24.69 | 22.18 | 15.64 to 28.71 | **** | <0.0001 |
| | *Ctrl vs. Emx1-cKO* | 46.87 | 27.08 | 19.78 | 13.25 to 26.32 | **** | <0.0001 |
| | *Nex-cKO vs. Emx1-cKO* | 24.69 | 27.08 | -2.393 | -9.555 to 4.769 | ns | 0.7036 |
| S | *Ctrl vs. Nex-cKO* | 23.25 | 27.56 | -4.312 | -10.85 to 2.226 | ns | 0.2608 |
| | *Ctrl vs. Emx1-cKO* | 23.25 | 23.42 | -0.1744 | -6.712 to 6.363 | ns | 0.9977 |
| | *Nex-cKO vs. Emx1-cKO* | 27.56 | 23.42 | 4.137 | -3.024 to 11.3 | ns | 0.3545 |
| A | *Ctrl vs. Nex-cKO* | 5.671 | 8.786 | -3.115 | -9.653 to 3.423 | ns | 0.4917 |
| | *Ctrl vs. Emx1-cKO* | 5.671 | 5.947 | -0.2756 | -6.813 to 6.262 | ns | 0.9944 |
| | *Nex-cKO vs. Emx1-cKO* | 8.786 | 5.947 | 2.839 | -4.323 to 10 | ns | 0.6105 |
| V | *Ctrl vs. Nex-cKO* | 10.13 | 21.5 | -11.36 | -17.9 to -4.825 | *** | 0.0003 |
| | *Ctrl vs. Emx1-cKO* | 10.13 | 21.35 | -11.22 | -17.76 to -4.681 | *** | 0.0003 |
| | *Nex-cKO vs. Emx1-cKO* | 21.5 | 21.35 | 0.1434 | -7.018 to 7.305 | ns | 0.9987 |
| RSC | *Ctrl vs. Nex-cKO* | 9.184 | 15.5 | -6.318 | -12.86 to 0.2199 | ns | 0.0604 |
| | *Ctrl vs. Emx1-cKO* | 9.184 | 19.01 | -9.821 | -16.36 to -3.284 | ** | 0.0017 |
| | *Nex-cKO vs. Emx1-cKO* | 15.5 | 19.01 | -3.503 | -10.67 to 3.658 | ns | 0.4734 |

**Figure 3E– Cortical distribution of YFP-H positive cells (#of cells/slides)**

| Area | Hypothesis | Mean 1 | Mean 2 | Mean diff. | 95,00% CI of diff | Summary | Adjusted P. |
|---|---|---|---|---|---|---|---|
| PFC | *Ctrl vs. Nex-cKO* | *33.06* | *6.42* | *26.64* | *-52.67 to 105.9* | *ns* | *0.5048* |
| | *Ctrl vs. Emx1-cKO* | *33.06* | *40.33* | *-7.269* | *-86.57 to 72.03* | *ns* | *0.8553* |
| | *Nex-cKO vs. Emx1-cKO* | *6.42* | *40.33* | *-33.91* | *-120.8 to 52.96* | *ns* | *0.4386* |
| M | *Ctrl vs. Nex-cKO* | *324.5* | *91.87* | *232.6* | *153.3 to 311.9* | ****** | *<0.0001* |
| | *Ctrl vs. Emx1-cKO* | *324.5* | *301.7* | *22.78* | *-56.52 to 102.1* | *ns* | *0.5682* |
| | *Nex-cKO vs. Emx1-cKO* | *91.87* | *301.7* | *-209.9* | *-296.7 to -123* | ****** | *<0.0001* |
| S | *Ctrl vs. Nex-cKO* | *163.2* | *105.2* | *57.98* | *-21.32 to 137.3* | *ns* | *0.1491* |
| | *Ctrl vs. Emx1-cKO* | *163.2* | *257.8* | *-94.58* | *-173.9 to -15.28* | *** | *0.0202* |
| | *Nex-cKO vs. Emx1-cKO* | *105.2* | *257.8* | *-152.6* | *-239.4 to -65.69* | ***** | *0.0008* |
| A | *Ctrl vs. Nex-cKO* | *38.09* | *32.77* | *5.32* | *-73.98 to 84.62* | *ns* | *0.8939* |
| | *Ctrl vs. Emx1-cKO* | *38.09* | *61.63* | *-23.53* | *-102.8 to 55.77* | *ns* | *0.5555* |
| | *Nex-cKO vs. Emx1-cKO* | *32.77* | *61.63* | *-28.85* | *-115.7 to 58.02* | *ns* | *0.5095* |
| V | *Ctrl vs. Nex-cKO* | *81.24* | *81.07* | *0.1681* | *-79.13 to 79.47* | *ns* | *0.9966* |
| | *Ctrl vs. Emx1-cKO* | *81.24* | *228* | *-146.8* | *-226.1 to -67.45* | ***** | *0.0004* |
| | *Nex-cKO vs. Emx1-cKO* | *81.07* | *228* | *-146.9* | *-233.8 to -60.05* | **** | *0.0012* |
| RSC | *Ctrl vs. Nex-cKO* | *64.39* | *58.41* | *5.974* | *-73.33 to 85.28* | *ns* | *0.8809* |
| | *Ctrl vs. Emx1-cKO* | *64.39* | *204.1* | *-139.8* | *-219.1 to -60.46* | ***** | *0.0008* |
| | *Nex-cKO vs. Emx1-cKO* | *58.41* | *204.1* | *-145.7* | *-232.6 to -58.86* | **** | *0.0013* |

**Figure 3F– Distribution of YFP-H positive cells among M and S areas (% values)**

| Area | Hypothesis | Mean 1 | Mean 2 | Mean diff. | 95,00% CI of diff | Summary | Adjusted P. |
|---|---|---|---|---|---|---|---|
| M | *Ctrl vs. Nex-cKO* | *66.85* | *47.35* | *19.5* | *7.864 to 31.14* | **** | *0.0021* |
| | *Ctrl vs. Emx1-cKO* | *66.85* | *53.43* | *13.43* | *1.79 to 25.07* | *** | *0.0257* |
| | *Nex-cKO vs. Emx1-cKO* | *47.35* | *53.43* | *-6.074* | *-18.82 to 6.676* | *ns* | *0.3339* |

| | | | | | | | |
|---|---|---|---|---|---|---|---|
| S | *Ctrl vs. Nex-cKO* | *33.15* | *52.65* | *-19.5* | *-31.14 to -7.864* | **** | *0.0021* |
| | *Ctrl vs. Emx1-cKO* | *33.15* | *46.57* | *-13.43* | *-25.07 to -1.79* | *** | *0.0257* |
| | *Nex-cKO vs. Emx1-cKO* | *52.65* | *46.57* | *6.074* | *-6.676 to 18.82* | *ns* | *0.3339* |

**Figure 4E – Rostral LFP diameter**

| Section | Hypothesis | Mean 1 | Mean 2 | Mean diff. | 95,00% CI of diff | Summary | Adjusted P. |
|---|---|---|---|---|---|---|---|
| 1 | *Ctrl vs. Nex-cKO* | 0 | 0 | 0 | -116.9 to 116.9 | ns | >0.9999 |
| | *Ctrl vs. Emx1-cKO* | 0 | 0 | 0 | -116.9 to 116.9 | ns | >0.9999 |
| | *Nex-cKO vs. Emx1-cKO* | 0 | 0 | 0 | -124.9 to 124.9 | ns | >0.9999 |
| 2 | *Ctrl vs. Nex-cKO* | 0 | 28.29 | -28.29 | -145.2 to 88.57 | ns | 0.836 |
| | *Ctrl vs. Emx1-cKO* | 0 | 0 | 0 | -116.9 to 116.9 | ns | >0.9999 |
| | *Nex-cKO vs. Emx1-cKO* | 28.29 | 0 | 28.29 | -96.64 to 153.2 | ns | 0.8549 |
| 3 | *Ctrl vs. Nex-cKO* | 0 | 132.7 | -132.7 | -249.6 to -15.84 | * | 0.0215 |
| | *Ctrl vs. Emx1-cKO* | 0 | 0 | 0 | -116.9 to 116.9 | ns | >0.9999 |
| | *Nex-cKO vs. Emx1-cKO* | 132.7 | 0 | 132.7 | 7.775 to 257.6 | * | 0.0343 |
| 4 | *Ctrl vs. Nex-cKO* | 0 | 179.6 | -179.6 | -296.5 to -62.77 | ** | 0.001 |
| | *Ctrl vs. Emx1-cKO* | 0 | 0 | 0 | -116.9 to 116.9 | ns | >0.9999 |
| | *Nex-cKO vs. Emx1-cKO* | 179.6 | 0 | 179.6 | 54.71 to 304.6 | ** | 0.0023 |
| 5 | *Ctrl vs. Nex-cKO* | 0 | 175.9 | -175.9 | -292.8 to -59.04 | ** | 0.0013 |
| | *Ctrl vs. Emx1-cKO* | 0 | 55.34 | -55.34 | -172.2 to 61.52 | ns | 0.5052 |
| | *Nex-cKO vs. Emx1-cKO* | 175.9 | 55.34 | 120.6 | -4.369 to 245.5 | ns | 0.0612 |
| 6 | *Ctrl vs. Nex-cKO* | 24.35 | 169.6 | -145.2 | -262.1 to -28.37 | * | 0.0103 |
| | *Ctrl vs. Emx1-cKO* | 24.35 | 115 | -90.67 | -207.5 to 26.2 | ns | 0.1624 |
| | *Nex-cKO vs. Emx1-cKO* | 169.6 | 115 | 54.57 | -70.37 to 179.5 | ns | 0.5592 |
| 7 | *Ctrl vs. Nex-cKO* | 117 | 231.4 | -114.4 | -231.3 to 2.467 | ns | 0.0565 |
| | *Ctrl vs. Emx1-cKO* | 117 | 234.6 | -117.7 | -234.5 to -0.8201 | * | 0.048 |
| | *Nex-cKO vs. Emx1-cKO* | 231.4 | 234.6 | -3.287 | -128.2 to 121.6 | ns | 0.9979 |
| 8 | *Ctrl vs. Nex-cKO* | 241.7 | 187.5 | 54.13 | -66.26 to 174.5 | ns | 0.5401 |
| | *Ctrl vs. Emx1-cKO* | 241.7 | 293 | -51.33 | -171.7 to 69.06 | ns | 0.5746 |
| | *Nex-cKO vs. Emx1-cKO* | 187.5 | 293 | -105.5 | -230.4 to 19.47 | ns | 0.1168 |
| 9 | *Ctrl vs. Nex-cKO* | 316.6 | 267.9 | 48.71 | -68.15 to 165.6 | ns | 0.5888 |
| | *Ctrl vs. Emx1-cKO* | 316.6 | 250.1 | 66.55 | -56.81 to 189.9 | ns | 0.4128 |
| | *Nex-cKO vs. Emx1-cKO* | 267.9 | 250.1 | 17.83 | -113.2 to 148.9 | ns | 0.9449 |
| 10 | *Ctrl vs. Nex-cKO* | 318.9 | 194.4 | 124.6 | 7.715 to 241.4 | * | 0.0335 |
| | *Ctrl vs. Emx1-cKO* | 318.9 | 205.8 | 113.1 | -3.738 to 230 | ns | 0.0602 |
| | *Nex-cKO vs. Emx1-cKO* | 194.4 | 205.8 | -11.45 | -136.4 to 113.5 | ns | 0.9746 |
| 11 | *Ctrl vs. Nex-cKO* | 231.1 | 88.06 | 143.1 | -163 to 449.1 | ns | 0.514 |
| | *Ctrl vs. Emx1-cKO* | 231.1 | 64.04 | 167.1 | -82.79 to 416.9 | ns | 0.258 |
| | *Nex-cKO vs. Emx1-cKO* | 88.06 | 64.04 | 24.02 | -225.8 to 273.9 | ns | 0.9721 |
| 12 | *Ctrl vs. Nex-cKO* | 276.1 | 179.6 | 96.5 | -26.86 to 219.9 | ns | 0.1576 |
| | *Ctrl vs. Emx1-cKO* | 276.1 | 136.5 | 139.6 | 16.23 to 263 | * | 0.022 |
| | *Nex-cKO vs. Emx1-cKO* | 179.6 | 136.5 | 43.09 | -93.77 to 179.9 | ns | 0.7389 |
| 13 | *Ctrl vs. Nex-cKO* | 201.9 | 180.5 | 21.33 | -243.7 to 286.3 | ns | 0.9804 |
| | *Ctrl vs. Emx1-cKO* | 201.9 | 44.45 | 157.4 | -40.14 to 354.9 | ns | 0.1471 |
| | *Nex-cKO vs. Emx1-cKO* | 180.5 | 44.45 | 136.1 | -113.8 to 385.9 | ns | 0.4059 |
| 14 | *Ctrl vs. Nex-cKO* | 218.4 | 114.6 | 103.8 | -19.52 to 227.2 | ns | 0.1182 |
| | *Ctrl vs. Emx1-cKO* | 218.4 | 95.41 | 123 | -0.3344 to 246.4 | ns | 0.0508 |
| | *Nex-cKO vs. Emx1-cKO* | 114.6 | 95.41 | 19.19 | -117.7 to 156 | ns | 0.9416 |
| 15 | *Ctrl vs. Nex-cKO* | 154.7 | 59.11 | 95.61 | -21.25 to 212.5 | ns | 0.1328 |
| | *Ctrl vs. Emx1-cKO* | 154.7 | 51.17 | 103.6 | -13.31 to 220.4 | ns | 0.0942 |
| | *Nex-cKO vs. Emx1-cKO* | 59.11 | 51.17 | 7.94 | -117 to 132.9 | ns | 0.9877 |
| 16 | *Ctrl vs. Nex-cKO* | 101.6 | 16.85 | 84.78 | -32.09 to 201.6 | ns | 0.2035 |
| | *Ctrl vs. Emx1-cKO* | 101.6 | 18.75 | 82.87 | -33.99 to 199.7 | ns | 0.2183 |
| | *Nex-cKO vs. Emx1-cKO* | 16.85 | 18.75 | -1.908 | -126.8 to 123 | ns | 0.9993 |
| 17 | *Ctrl vs. Nex-cKO* | 60.94 | 0 | 60.94 | -55.93 to 177.8 | ns | 0.4374 |
| | *Ctrl vs. Emx1-cKO* | 60.94 | 0 | 60.94 | -55.93 to 177.8 | ns | 0.4374 |
| | *Nex-cKO vs. Emx1-cKO* | 0 | 0 | 0 | -124.9 to 124.9 | ns | >0.9999 |
| | *Ctrl vs. Nex-cKO* | 28.85 | 0 | 28.85 | -88.02 to 145.7 | ns | 0.8301 |

| | | | | | | | |
|---|---|---|---|---|---|---|---|
| **18** | *Ctrl vs. Emx1-cKO* | 28.85 | 0 | 28.85 | -88.02 to 145.7 | ns | 0.8301 |
| | *Nex-cKO vs. Emx1-cKO* | 0 | 0 | 0 | -124.9 to 124.9 | ns | >0.9999 |
| **19** | *Ctrl vs. Nex-cKO* | 0 | 0 | 0 | -112 to 112 | ns | >0.9999 |
| | *Ctrl vs. Emx1-cKO* | 0 | 0 | 0 | -116.9 to 116.9 | ns | >0.9999 |
| | *Nex-cKO vs. Emx1-cKO* | 0 | 0 | 0 | -120.4 to 120.4 | ns | >0.9999 |

**Figure 4F – Caudal LFP diameter**

| Section | Hypothesis | Mean 1 | Mean 2 | Mean diff. | 95,00% CI of diff | Summary | Adjusted P. |
|---|---|---|---|---|---|---|---|
| 1 | *Ctrl vs. Nex-cKO* | 0 | 0 | 0 | -88.51 to 88.51 | ns | >0.9999 |
| | *Ctrl vs. Emx1-cKO* | 0 | 0 | 0 | -88.51 to 88.51 | ns | >0.9999 |
| | *Nex-cKO vs. Emx1-cKO* | 0 | 0 | 0 | -94.62 to 94.62 | ns | >0.9999 |
| 2 | *Ctrl vs. Nex-cKO* | 0 | 46.89 | -46.89 | -135.4 to 41.62 | ns | 0.4252 |
| | *Ctrl vs. Emx1-cKO* | 0 | 0 | 0 | -88.51 to 88.51 | ns | >0.9999 |
| | *Nex-cKO vs. Emx1-cKO* | 46.89 | 0 | 46.89 | -47.73 to 141.5 | ns | 0.4728 |
| 3 | *Ctrl vs. Nex-cKO* | 0 | 98.1 | -98.1 | -186.6 to -9.584 | * | 0.0257 |
| | *Ctrl vs. Emx1-cKO* | 0 | 0 | 0 | -88.51 to 88.51 | ns | >0.9999 |
| | *Nex-cKO vs. Emx1-cKO* | 98.1 | 0 | 98.1 | 3.473 to 192.7 | * | 0.0402 |
| 4 | *Ctrl vs. Nex-cKO* | 0 | 126.2 | -126.2 | -214.7 to -37.72 | ** | 0.0026 |
| | *Ctrl vs. Emx1-cKO* | 0 | 0 | 0 | -88.51 to 88.51 | ns | >0.9999 |
| | *Nex-cKO vs. Emx1-cKO* | 126.2 | 0 | 126.2 | 31.6 to 220.9 | ** | 0.0053 |
| 5 | *Ctrl vs. Nex-cKO* | 18.5 | 111 | -92.51 | -181 to -3.999 | * | 0.0382 |
| | *Ctrl vs. Emx1-cKO* | 18.5 | 13.31 | 5.196 | -83.32 to 93.71 | ns | 0.9895 |
| | *Nex-cKO vs. Emx1-cKO* | 111 | 13.31 | 97.71 | 3.084 to 192.3 | * | 0.0412 |
| 6 | *Ctrl vs. Nex-cKO* | 69.28 | 128.1 | -58.82 | -147.3 to 29.69 | ns | 0.2617 |
| | *Ctrl vs. Emx1-cKO* | 69.28 | 97.47 | -28.19 | -116.7 to 60.32 | ns | 0.7331 |
| | *Nex-cKO vs. Emx1-cKO* | 128.1 | 97.47 | 30.63 | -63.99 to 125.3 | ns | 0.7256 |
| 7 | *Ctrl vs. Nex-cKO* | 107.5 | 176 | -68.56 | -300.3 to 163.2 | ns | 0.765 |
| | *Ctrl vs. Emx1-cKO* | 107.5 | 152.1 | -44.66 | -227.9 to 138.6 | ns | 0.8336 |
| | *Nex-cKO vs. Emx1-cKO* | 176 | 152.1 | 23.9 | -159.3 to 207.1 | ns | 0.9491 |
| 8 | *Ctrl vs. Nex-cKO* | 175.2 | 237.2 | -61.95 | -157.9 to 34.02 | ns | 0.2821 |
| | *Ctrl vs. Emx1-cKO* | 175.2 | 154.5 | 20.69 | -70.49 to 111.9 | ns | 0.8541 |
| | *Nex-cKO vs. Emx1-cKO* | 237.2 | 154.5 | 82.63 | -16.61 to 181.9 | ns | 0.1235 |
| 9 | *Ctrl vs. Nex-cKO* | 214.5 | 138.2 | 76.34 | -12.17 to 164.9 | ns | 0.1063 |
| | *Ctrl vs. Emx1-cKO* | 214.5 | 197.7 | 16.82 | -71.69 to 105.3 | ns | 0.8952 |
| | *Nex-cKO vs. Emx1-cKO* | 138.2 | 197.7 | -59.52 | -154.1 to 35.1 | ns | 0.3005 |
| 10 | *Ctrl vs. Nex-cKO* | 209.6 | 115.1 | 94.49 | 5.983 to 183 | * | 0.0333 |
| | *Ctrl vs. Emx1-cKO* | 209.6 | 132 | 77.6 | -10.91 to 166.1 | ns | 0.0988 |
| | *Nex-cKO vs. Emx1-cKO* | 115.1 | 132 | -16.89 | -111.5 to 77.73 | ns | 0.9069 |
| 11 | *Ctrl vs. Nex-cKO* | 232.5 | 99.13 | 133.4 | 44.9 to 221.9 | ** | 0.0013 |
| | *Ctrl vs. Emx1-cKO* | 232.5 | 116.2 | 116.4 | 27.87 to 204.9 | ** | 0.0061 |
| | *Nex-cKO vs. Emx1-cKO* | 99.13 | 116.2 | -17.03 | -111.7 to 77.59 | ns | 0.9055 |
| 12 | *Ctrl vs. Nex-cKO* | 186.4 | 127.8 | 58.65 | -34.78 to 152.1 | ns | 0.302 |
| | *Ctrl vs. Emx1-cKO* | 186.4 | 26.38 | 160.1 | 71.56 to 248.6 | **** | <0.0001 |
| | *Nex-cKO vs. Emx1-cKO* | 127.8 | 26.38 | 101.4 | 2.179 to 200.7 | * | 0.0439 |
| 13 | *Ctrl vs. Nex-cKO* | 94.47 | 0 | 94.47 | -137.3 to 326.2 | ns | 0.6019 |
| | *Ctrl vs. Emx1-cKO* | 94.47 | 13.38 | 81.09 | -95.93 to 258.1 | ns | 0.527 |
| | *Nex-cKO vs. Emx1-cKO* | 0 | 13.38 | -13.38 | -190.4 to 163.6 | ns | 0.9826 |
| 14 | *Ctrl vs. Nex-cKO* | 58.1 | 0 | 58.1 | -33.08 to 149.3 | ns | 0.2913 |
| | *Ctrl vs. Emx1-cKO* | 58.1 | 0 | 58.1 | -33.08 to 149.3 | ns | 0.2913 |
| | *Nex-cKO vs. Emx1-cKO* | 0 | 0 | 0 | -94.62 to 94.62 | ns | >0.9999 |
| 15 | *Ctrl vs. Nex-cKO* | 0 | 0 | 0 | -84.82 to 84.82 | ns | >0.9999 |
| | *Ctrl vs. Emx1-cKO* | 0 | 0 | 0 | -88.51 to 88.51 | ns | >0.9999 |
| | *Nex-cKO vs. Emx1-cKO* | 0 | 0 | 0 | -91.18 to 91.18 | ns | >0.9999 |

**Figure 4G – Rostral LFP area**

| | Hypothesis | Mean 1 | Mean 2 | Mean diff. | 95,00% CI of diff | Summary | Adjusted P. |
|---|---|---|---|---|---|---|---|

| | | | | | | | |
|---|---|---|---|---|---|---|---|
| | *Ctrl vs. Nex-cKO* | 2414 | 2258 | 156.4 | -570.3 to 883 | ns | 0.8257 |
| | *Ctrl vs. Emx1-cKO* | 2414 | 1725 | 689.2 | -37.42 to 1416 | ns | 0.0637 |
| **Figure 4H – Caudal LFP area** | | | | | | | |
| | **Hypothesis** | **Mean 1** | **Mean 2** | **Mean diff.** | **95,00% CI of diff** | **Summary** | **Adjusted P.** |
| | *Ctrl vs. Nex-cKO* | 1403 | 1427 | -24.29 | -261.8 to 213.2 | ns | 0.9566 |
| | *Ctrl vs. Emx1-cKO* | 1403 | 884.1 | 518.8 | 281.3 to 756.3 | *** | 0.0001 |
| **Figure 4I – Fasciculation Index** | | | | | | | |
| **Region** | **Hypothesis** | **Mean 1** | **Mean 2** | **Mean diff.** | **95,00% CI of diff** | **Summary** | **Adjusted P.** |
| **250 µm** | *Ctrl vs. Nex-cKO* | 0.6528 | 0.5493 | 0.1035 | 0.01185 to 0.1951 | * | 0.0216 |
| | *Ctrl vs. Emx1-cKO* | 0.6528 | 0.5595 | 0.09326 | 0.001658 to 0.1849 | * | 0.0446 |
| **500 µm** | *Ctrl vs. Nex-cKO* | 0.674 | 0.5771 | 0.0969 | 0.00254 to 0.1913 | * | 0.0422 |
| | *Ctrl vs. Emx1-cKO* | 0.674 | 0.5146 | 0.1594 | 0.06502 to 0.2537 | *** | 0.0004 |

***Supplementary Table 5: List of primary and secondary antibodies used in this study.***

| Antigen | Provider | Catalog # | Species | Working dilution |
|---|---|---|---|---|
| GFP | Abcam | Ab13970 | Ck | 1:500 |
| RFP | Abcam | Ab 124754 | Rb | 1:500 |
| Ck IgY - AF 488 | Thermo Fisher | A11039 | Gt | 1:500 |
| Rb IgG - AF 555 | Thermo Fisher | A21428 | Gt | 1:500 |
